# Peroxisomes regulate virulence and cell density sensing in *Cryptococcus neoformans*

**DOI:** 10.1101/2025.05.14.654083

**Authors:** Ella Jacobs, Quigly Dragotakes, Samuel Rodrigues dos Santos, Daniel Smith, Amanda Dziedzic, Anne Jedlicka, Barbara Smith, Julie M. Wolf, Carolina Coelho, Arturo Casadevall

## Abstract

*Cryptococcus neoformans*, a ubiquitous environmental fungus that causes cryptococcosis, survives in diverse environments including human hosts due to metabolic flexibility. Consequently, identifying how *C. neoformans* connects diverse metabolic pathways and virulence factor expression is important for understanding fungal pathogenesis. Peroxisomes play an essential role in metabolic homeostasis and regulation of carbon and lipid metabolism. In this article, we report a link between nickel exposure, a known hypoxia-mimetic and mitochondrial respiration inhibitor in yeast, and peroxisomal β-oxidation. Loss of the last two genes involved in the peroxisomal β-oxidation pathway, *MFE2* (CNAG_05721) and *POT1* (CNAG_00490), resulted in cell density-dependent virulence factor defects and growth inhibition attributed to a metabolic state involving large peroxisomes. We found that increasing cell density rescued virulence factor phenotypes and growth. Our results implicate mitochondrial retrograde signaling (RTG), a previously uncharacterized pathway in *C. neoformans*, in cell density sensing, peroxisomal β-oxidation pathway expression, and virulence, thus highlighting a critical role for metabolism in cryptococcal virulence.

## Introduction

Environmental microbes face unique nutritional challenges and diverse stressors [1, 2]. *Cryptococcus neoformans,* the causative agent of cryptococcosis, is a fungal pathogen with worldwide environmental distribution. *C. neoformans* is primarily found in bird excreta, soil, and trees [3]. Most severe cases of cryptococcosis occur in immunocompromised individuals, where the most common presentation is meningitis, which is fatal if untreated [4].

*C. neoformans* senses and responds to environmental cues with metabolic plasticity, which allows the fungal pathogen to survive in diverse environments and multiple hosts. As evidence of this plasticity, phagocytosis induces immediate metabolic reprogramming including pathways related to the tricarboxylic acid cycle (TCA cycle), glyoxylate pathway, gluconeogenesis, lipid catabolism and two-carbon metabolism [5–9]. The upstream regulatory networks that control the convergence of the phagocytosis-induced pathways are unknown [10]. Similarly to the host environment, *C. neoformans* senses and responds to fungal cell-density with a quorum-sensing response integrated into nutrient acquisition, mating, virulence, and metabolism [11–14]. This process is controlled by transcription factors Qsp1, Nrg1, Liv3, and Cqs2 [11–14]. In contrast, far less is known about how *C. neoformans* senses low cell density. The fungal response to low cell density is important as it mirrors how the fungal cells enter the host in low numbers often within macrophage phagosomes.

Mitochondria-mediated stress responses are of particular interest in *C. neoformans* because functional mitochondria are essential for virulence [15–17]. Nickel exposure induces high levels of oxidative stress that inhibit mitochondrial function [18]. Given the importance of the mitochondria as a metabolic signaling hub and thus for coordinating metabolism during infection, we wondered if disrupting respiration using nickel would reveal signaling pathways responsible for integrating infection-induced pathways. Here, we analyzed gene expression upon exposure of *C. neoformans* cells to subinhibitory concentrations of nickel during starvation in minimal media. This revealed induction of a metabolic program involving peroxisomal β-oxidation genes and peroxisomal biogenesis. In *C. neoformans,* a mechanistic link between mitochondria and peroxisomes was previously uncharacterized. Within the peroxisome, β-oxidation converts fatty acyl-CoA into acetyl-CoA via the enzymes Faa2 (encoded by CNAG_03019), Pox1 (encoded by CNAG_07747), Mfe2 (also Fox2, encoded by CNAG_05721), and Pot1 (encoded by CNAG_00490).

Previously, Kretschmer et al. demonstrated that disruption of the cryptococcal *MFE2* gene resulted in diverse phenotypes including virulence factor defects, fluconazole susceptibility, and sensitivity to cell wall stressors [19]. Interestingly, they also uncovered a growth defect of the *mfe2*Δ strain. The growth inhibition of *mfe2*Δ was rescued by an unidentified secreted factor produced by both wild-type and *qsp1*Δ fungal cells and bacterial colonies of *E. coli* and *S. aureus* [19]. The cause of such pleotropic phenotypes was attributed to metabolic perturbations in diverse pathways including nitrogen, lipid, and carbon metabolism [19]. Here, we show that mitochondrial retrograde signaling (RTG), an uncharacterized pathway in *C. neoformans*, is the regulator of this cell-density-dependent metabolic state.

Here, we show that disruption of *MFE2* and *POT1,* the genes encoding enzymes which collectively perform the three final steps of peroxisomal β-oxidation, resulted in cell-density-dependent growth and virulence factor defects. By increasing cell density, we rescued the growth and virulence factor defects of *mfe2*Δ and *pot1*Δ, illustrating that the mutant strains produce the secreted factor required for growth rescue. Furthermore, we implicate RTG signaling in cell-density sensing and the metabolic response to infection.

## Results

### Nickel exposure induces peroxisomal gene expression in *C. neoformans*

Nickel is a known hypoxia mimetic and mitochondrial respiration inhibitor in eukaryotes [18, 20–23]. We were interested in describing the stress response to simultaneous nickel exposure and starvation with the goal of identifying mitochondrial-controlled stress pathways. Consequently, we cultured *C. neoformans* cells in minimal media with a sub-inhibitory concentration nickel sulfate (31.5 μM NiSO_4_) and performed gene expression analysis using bulk RNA-Seq.

Gene expression changes meeting the criteria of p < 0.05 and Fold-Change cutoff of -2 (blue) to 2 (red) were considered differentially expressed (Fig. 1A). Overall, 479 genes were differentially expressed after nickel exposure, of which 294 up-regulated and 185 down-regulated, compared to starvation alone (Supplementary Table 1). To characterize the response, we performed KEGG pathway analysis, solely for the up-regulated genes (Fig. 1B) and gene ontology (GO) term (Fig. 1C&D) for all the differentially expressed genes (FDR < 0.05). KEGG pathway analysis of the 294 genes downregulated during nickel exposure returned no pathways meeting the FDR < 0.05 cutoff.

**Figure 1:**
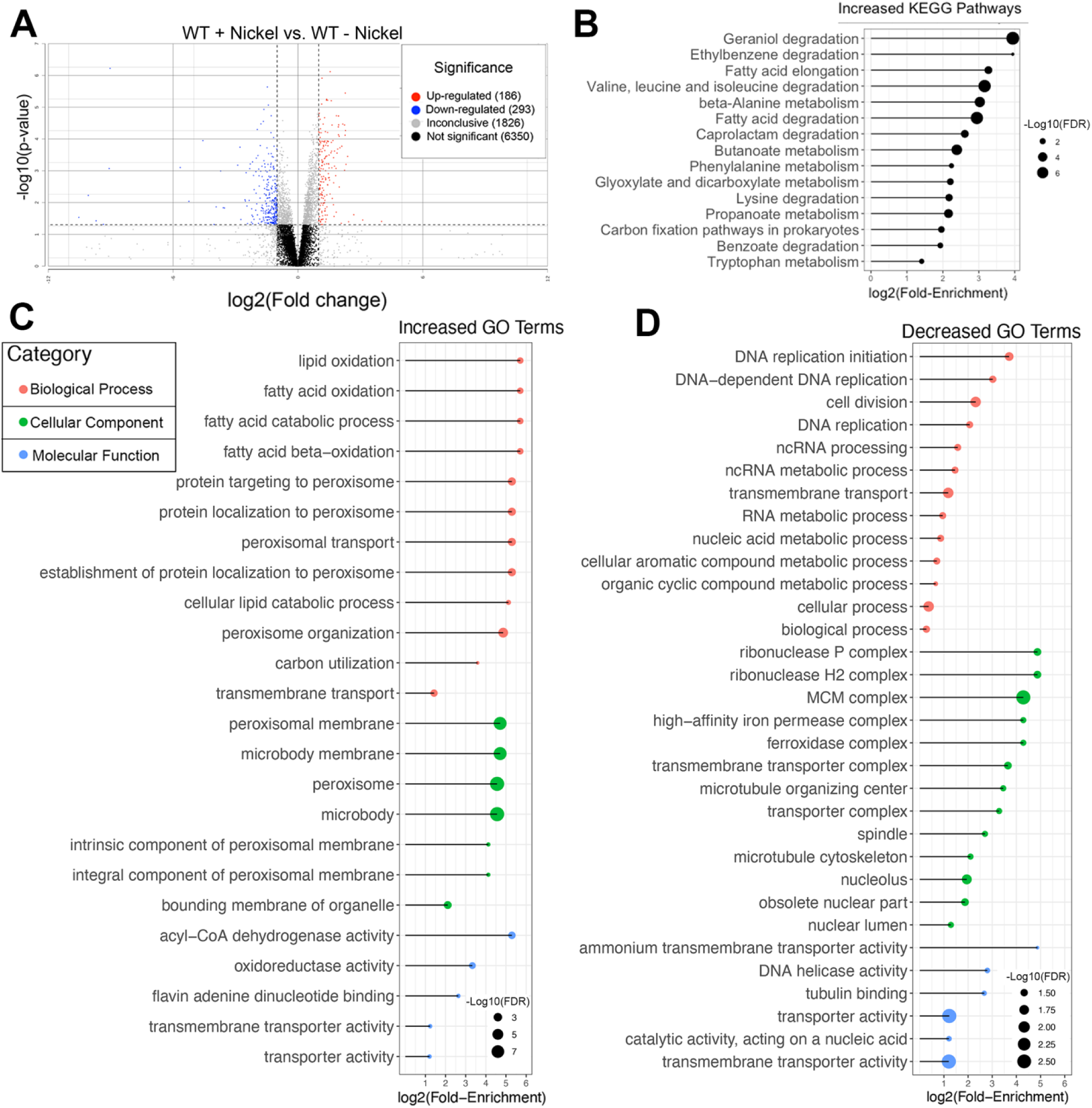
Nickel exposure induces peroxisomal gene expression in *C. neoformans*. **(A)** Volcano plot of H99 WT gene expression with and without NiSO_4_ (31.5 μM). Genes meeting the p-value < 0.05 and fold-change cutoff of -2 (blue) to +2 (red) were considered differentially expressed. **(B)** KEGG metabolic pathways increased during nickel exposure and starvation (FDR < 0.05 cutoff). No KEGG-pathways decreased during nickel exposure met this threshold. Circle size represents (−log_10_ [FDR]) and position represents (log_2_ [Fold-Enrichment]). **(C and D)** Gene ontology (GO) terms of differentially expressed genes (FDR < 0.05). GO processes **(C)** increased during nickel exposure and **(D)** decreased during nickel exposure. Colors represent GO term categories: Biological Processes (pink) Cellular Component (green) and Molecular Function (blue). Circle size represents (−log_10_ [FDR]) and position represents (log_2_ [Fold-Enrichment]).

Overall, we found that nickel exposure significantly increased several metabolic pathways and GO term categories related to lipid catabolism and peroxisomes (Fig.1B-D). This is consistent with the recently characterized role of *C. neoformans* fatty acid catabolism and peroxisomal biogenesis in survival during long-term starvation and hypoxia [24]. Additionally, the degradation pathways of amino acids (valine, leucine, isoleucine, and lysine), the degradation of carboxylic acids (propanoate, butanoate, and benzoate), and the metabolism of amino acids (tryptophan, phenylalanine, and β-alanine) were increased after nickel exposure. This is consistent with *S. cerevisiae* metabolome analysis demonstrating that amino acid metabolism is dramatically impacted by hypoxia [22].

### Peroxisomal β-oxidation genes affect cryptococcal stress tolerance, virulence factors, and growth

Nickel exposure induces high levels of oxidative stress that inhibit mitochondrial function [18]. We hypothesized that a metabolic shift occurred to compensate for inhibited mitochondrial function. Given that a direct link between the mitochondria and peroxisomal lipid metabolism was unexplored in *C. neoformans*, we focused on the peroxisomal β-oxidation pathway for further characterization (Fig. 2A). Long-chain fatty acids (LCFA) and very-long-chain fatty acids (VLCFA) are converted to activated fatty acyl-CoA by cytosolic FAA1-FAA4, followed by peroxisomal import via Pxa1p (peroxisomal ABC-transporter 1) and Pxa2p [25–27]. Medium-chain fatty acids (MCFA) diffuse freely across the peroxisomal membrane and are subsequently activated by acyl-CoA synthetase Faa2 (encoded by CNAG_03019) [28]. The first step of β-oxidation is performed in a rate-limiting manner by peroxisomal acyl-CoA oxidase Pox1 (encoded by CNAG_07747) [29]. The multi-functional enzyme Mfe2, (also Fox2, encoded by CNAG_05721) then performs two functions: firstly, as a 2-enoyl-CoA hydratase 2 and secondly as a D-3-hydroxyacyl CoA dehydrogenase to produce 3-ketoacyl-CoA [30]. Finally, Pot1 (encoded by CNAG_00490), a 3-ketoacyl-CoA thiolase, cleaves the 3-ketoacyl-CoA intermediate to yield one acetyl-CoA and an acyl-CoA shortened by 2 carbons [31]. This process repeats several times to fully oxidize long-chain fatty acids.

**Figure 2:**
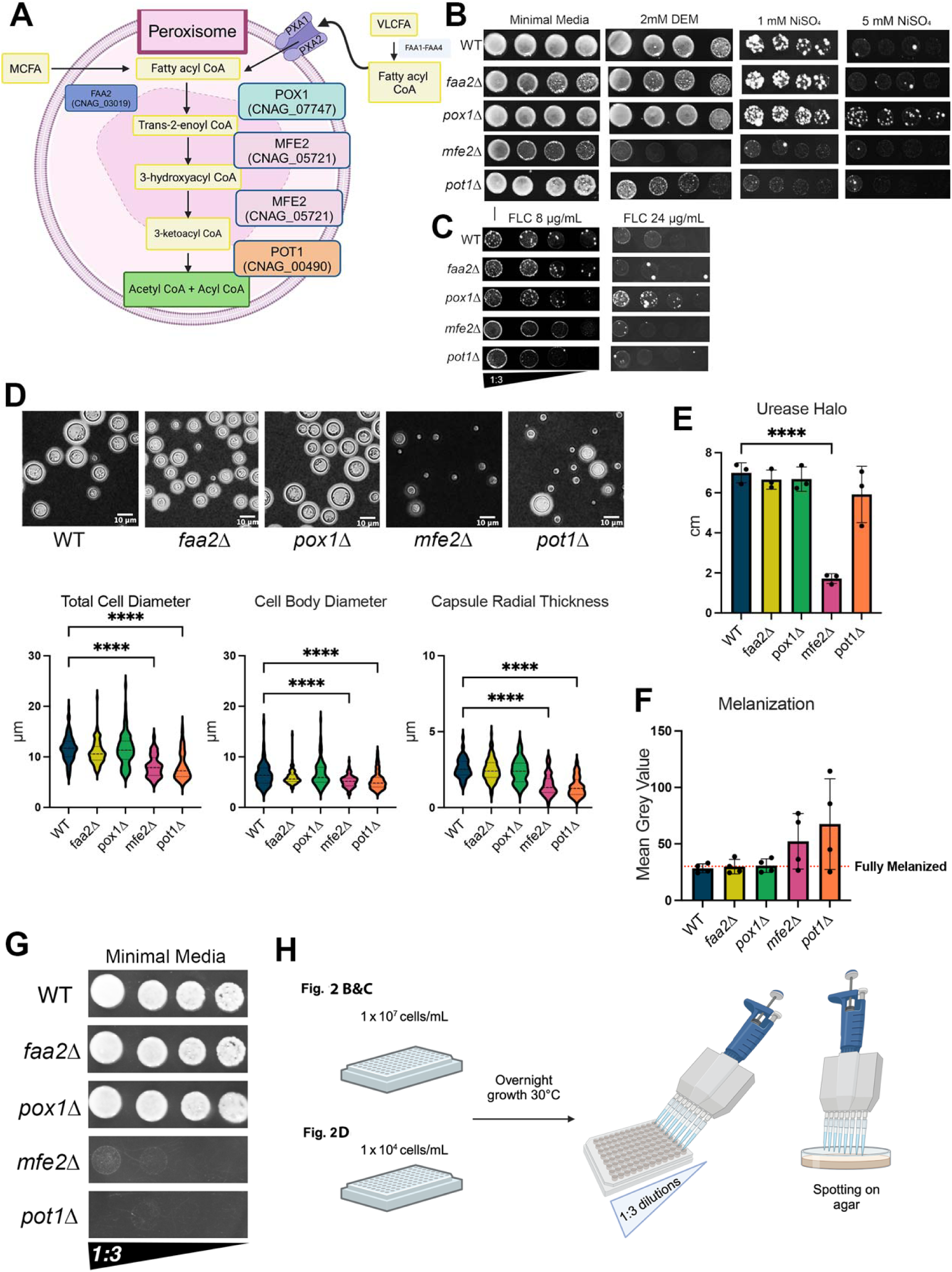
Peroxisomal β-oxidation genes affect cryptococcal stress tolerance, virulence factors and growth. **(A)** Peroxisomal β-oxidation pathway of *Saccharomyces cerevisiae* with *C. neoformans* homologs (CNAGs). Created with BioRender. **(B)** *mfe2*Δ and *pot1*Δ were sensitive to the oxidative stress inducer Diethyl Maleate (DEM). Strains *mfe2*Δ and *pot1*Δ were sensitive to nickel exposure while *pox1*Δ had high nickel tolerance, which was especially apparent at 5 mM. 1:3 serial dilutions were spotted on minimal media agar containing either DEM (2mM) or nickel and imaged after 48 h of growth at 30°C. *pox1*Δ had high nickel tolerance, which was especially apparent at 5 mM. **(C)** Susceptibility to fluconazole exposure. At 8 μg/mL, *mfe2*Δ and *pot1*Δ displayed similar fluconazole tolerance to WT at higher cell density spots and reduced tolerance in lower cell density spots. At 24 μg/mL of fluconazole, only *pox1*Δ was tolerant. The colony photographs throughout the paper were taken under the same illumination and consequently some colonies appear to show very similar or identical light reflections that manifest as white dots, arcs, and other light effects. However, these photographs were each individually taken and were not spliced or combined, thus represent different sets of colonies grown under different conditions. **D)** Representative India ink images after 7 d in minimal media without L-DOPA (40x magnification, scale bar = 10 μm). Measurements of cell diameter (capsule + cell body), cell body diameter, and capsule radial thickness. Compared to WT, the *mfe2*Δ and *pot1*Δ strains had significantly smaller cell body sizes and capsule radial thickness (n= 3 biological replicates, ∼25 cells measured and pooled between biological replicates, One-way ANOVA ****, p-value <0.0001). **E)** Urease halo size (n = 3 biological replicates after 48 h). *mfe2*Δ had a severe urease defect compared to WT (One-way ANOVA, p-value <0.0001), and *pot1*Δ produced urease halos with ∼3-times the standard deviation of WT (WT: Mean=6.998 SD=0.4988, *mfe2*Δ: Mean= 1.716 SD=0.2490, *pot1*Δ: Mean=5.914 SD=1.406). **F)** Mean grey value of melanization cultures showed highly variable growth and melanization for both *mfe2*Δ and *pot1*Δ (n=4 biological replicates, WT: Mean=28.35, SD= 3.940, *mfe2*Δ: Mean=52.24 SD= 24.59, *pot1*Δ: Mean=67.52 SD=40.16). **G)** Low starting inoculum (1 x 10^4^ cells/mL) arrests overnight growth of *mfe2*Δ and *pot1*Δ, resulting in reduced CFUs. **H)** Starting inoculum differed between serial dilution experiments in figure 2 panels B & C (1 x 10^7^ cells/mL) and figure 2 panel G (1 x 10^4^ cells/mL). Decreasing the starting inoculum markedly reduced the number of CFUs of both mutants compared to WT.

We reasoned that the deletion of the compensatory peroxisomal β-oxidation pathway should increase sensitivity to oxidative stress induced by nickel and by other oxidative stressors. We spotted strains on minimal media agar containing nickel (1 mM and 5 mM NiSO_4_) (Fig. 2B). *mfe2*Δ and *pot1*Δ were sensitive to nickel exposure while *pox1*Δ was tolerant. Exposure to oxidative stress inducer Diethyl Maleate (DEM) revealed *mfe2*Δ and *pot1*Δ were sensitive to DEM-induced oxidative stress (Fig. 2B).

Mitochondrial dysfunction results in azole drug resistance in *C. neoformans* [16], and *mfe2*Δ was previously demonstrated to have fluconazole susceptibility [19]. To assess fluconazole (FLC) tolerance, strains were exposed 8 μg/mL (the established minimum inhibitory concentration (MIC) [32, 33]) and 24 μg/mL (Fig. 2C). At 8 μg/mL, *mfe2*Δ and *pot1*Δ displayed similar fluconazole tolerance to WT (KN99α) at high cell density and with reduced tolerance in low cell density spots. Interestingly, the cell-density dependent growth differences are also apparent in the known stress phenotypes of *mfe2*Δ [19]. At 24 μg/mL, *mfe2*Δ and *pot1*Δ had similar tolerance as WT, while *pox1*Δ was fluconazole resistant. Taken together, these results revealed *mfe2*Δ and *pot1*Δ are sensitive to nickel, DEM, and fluconazole, implicating the peroxisomal pathway as an important compensatory pathway during mitochondrial stress.

*C. neoformans* uses several virulence factors to survive in the host and cause disease, including urease secretion, melanization, and the production of a polysaccharide capsule. The *mfe2*Δ strain has previously been characterized for defects in capsule and melanin production [19]. In agreement with these results, when we assessed virulence factor expression in the context of peroxisomal function, we similarly found defects in virulence factors for *mfe2*Δ and *pot1*Δ. However, these phenotypes were accompanied by a high level of variability between biological replicates.

To assess capsule production, we visualized the capsule with India ink negative staining (Fig. 2D) and measured the total cell diameter, cell body diameter, and capsule radial thickness. Visually, *mfe2*Δ and *pot1*Δ were a mix of small cells lacking capsules and larger fungal cells with capsules. Compared to WT, *mfe2*Δ and *pot1*Δ had significantly smaller cell body sizes and capsule radial thickness (∼25 cells measured per biological triplicate, One-way ANOVA ****, p-value <0.0001). To test urease secretion, we spotted strains on Christensen’s urea agar and measured the color halo resulting from the ammonia mediated pH change (Fig. 2E). Compared to WT, *mfe2*Δ had a severe urease defect (n=3 biological replicates, one-way ANOVA, p-value <0.0001) and *pot1*Δ produced urease halos with ∼3-times the standard deviation of WT (WT: Mean(±SD):= 6.998±0.4988, *mfe2*Δ: Mean(±SD):= 1.716±0.2490, *pot1*Δ: Mean(±SD):= 5.914, ±1.406). We then assessed cellular melanization in minimal media supplemented with 1 mM L-DOPA (Fig. 2F). Melanization was measured as the mean grey value of cell cultures, where a lower mean grey value indicates a highly melanized cell culture. Phenotypic variability was also apparent when comparing cellular melanization: compared to WT, *mfe2*Δ and *pot1*Δ had large standard deviations (n=4 biological replicates, one-way ANOVA, WT: Mean(±SD)= 28.35 (±3.940), *mfe2*Δ: Mean(±SD)= 52.24 (± 24.59), *pot1*Δ: Mean(±SD)= 67.52 (±40.16).

Given the high variability of the virulence factor results and the cell-density-dependent growth in response to stress, we considered that a cell-density-dependent growth defect was responsible for the virulence factor variability. We hypothesized that starting a culture with a low initial inoculum and allowing overnight growth would arrest the growth of the mutants, resulting in reduced CFUs. To test this, we lowered the starting inoculum used in Figure 2 panels B & C (Fig. 2G&H). Lowering the starting inoculum markedly reduced the CFUs of both *mfe2*Δ and *pot1*Δ (Fig. 2G). Taken together, these data show that both *mfe2*Δ and *pot1*Δ mutant strains had highly variable defects in key cryptococcal virulence factors. We found that both strains had growth defects at low starting inocula and hypothesized that the variability we observed was due to this growth defect. Additionally, we screened 4 other strains with knockouts in genes upregulated during nickel exposure for responses to relevant stresses and urease secretion and found that only *mfe2*Δ and *pot1*Δ exhibited nickel sensitivity (Sup. Fig. 1).

### Commonalities in the transcriptional response to nickel exposure, pigeon guano, stationary phase, and high-density

We hypothesized that nickel exposure induced RTG pathway expression to compensate for inhibited mitochondrial function using peroxisomal genes and other anaplerotic pathways [34]. The details of mitochondrial retrograde signaling pathways are not conserved between eukaryotes [35]. *C. neoformans* lacks known homologs of *RTG1* and *RTG2* [36], but does possess an *RTG3* homolog known as *MLN1* [36, 37].

Our initial experiments hinted that the RTG-associated pathways induced upon nickel exposure are also regulated by cell density and growth stage. To probe this, we compared the genes differentially expressed in our nickel-exposed mRNA-Seq dataset to 3 previously published mRNA-Seq datasets comparing *C. neoformans* gene expression in stationary phase vs. log phase [24] and 2.5 OD vs. log phase [13]. We also compared our nickel-exposed mRNA-Seq dataset to pigeon guano vs. YPD [38], since in *S. cerevisiae,* nitrogen sources ammonia and urea activate RTG signaling [39]. Across all 4 mRNA-Seq datasets, we found an overlap of 98 differentially expressed genes (Fig. 3A and B).

**Figure 3:**
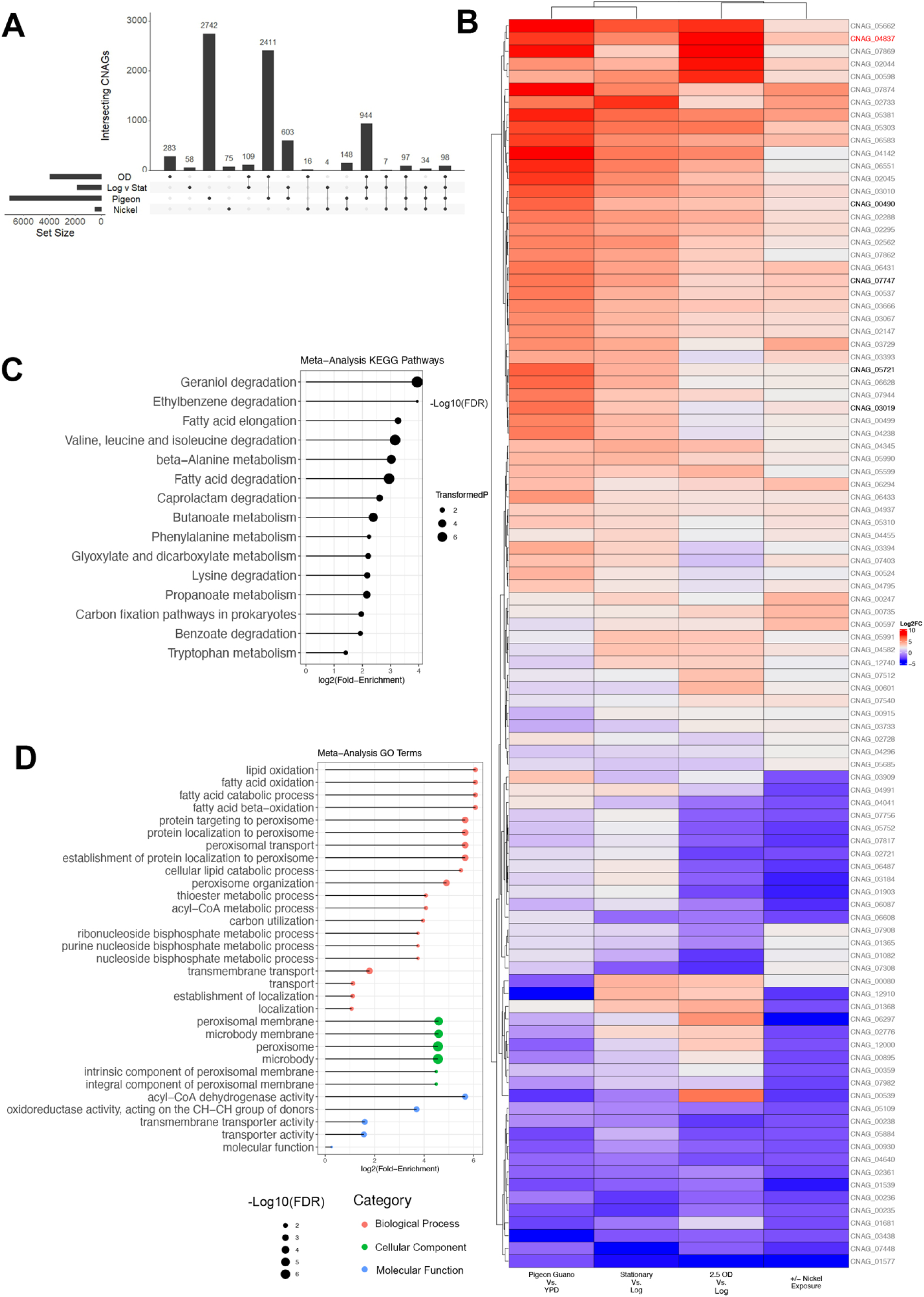
Commonalities in the transcriptional response to nickel exposure, pigeon guano, stationary phase, and high-OD. **(A)** Intersecting differentially expressed genes across RNAseq datasets. Comparing significantly differentially expressed genes between nickel exposure, stationary vs. log phase, 2.5 OD vs. log phase, and pigeon guano vs. YPD. 98 genes were differentially expressed in all 4 datasets. **(B)** Heatmap showing fold-change differences of the 98 genes within each dataset. Peroxisomal β-oxidation pathway genes (bolded) and *MLN1* (red). **(C)** KEGG metabolic pathways of the 98 genes differentially expressed in all 4 datasets (FDR < 0.05 cutoff). Circle size represents (−log_10_ [FDR]) and position represents (log_2_ [Fold-Enrichment]). **(D)** Gene ontology (GO) terms of the 98 meta-analysis identified genes (FDR < 0.05). Colors represent GO term categories: Biological Processes (pink) Cellular Component (green) and Molecular Function (blue). Circle size represents (−log_10_ [FDR]) and position represents (log_2_ [Fold-Enrichment]).

The peroxisomal β-oxidation pathway genes (bolded) and *MLN1* (red) were upregulated in the pigeon guano dataset, stationary phase relative to log-phase, and 2.5 OD relative to log-phase. We performed KEGG metabolic pathway analysis (Fig. 3C) and GO term analysis (Fig. 3D) on these 98 overlapping genes. Both KEGG analysis and GO term analysis returned similar pathways to our original analysis (Figure 1 panel B), indicating an RTG metabolic response is induced during nickel exposure and growth on pigeon guano, while being repressed during log-phase growth. We next sought to characterize this density-dependent growth defect.

### Cell-density-related effects on growth

In Figure 2 Panel G, we demonstrated that lowering the pre-culture cell density inhibited growth. For future experiments, we wanted to avoid pre-stressing the culture with low cell density before experiments. Strains were streaked on minimal media agar to lawn density and resuspended in PBS in two cell densities: 1x10^7^ cells/mL and 1x10^6^ cells/mL. These two cell densities were serially diluted onto minimal media plates (Fig. 4A). The growth defects of *mfe2*Δ and *pot1*Δ were partially rescued at the higher cell density, but the growth defect was exacerbated at the lower cell density. This indicated that lower cell density could be sensed within individual agar spots and did not depend on low-density preculture conditions.

**Figure 4.**
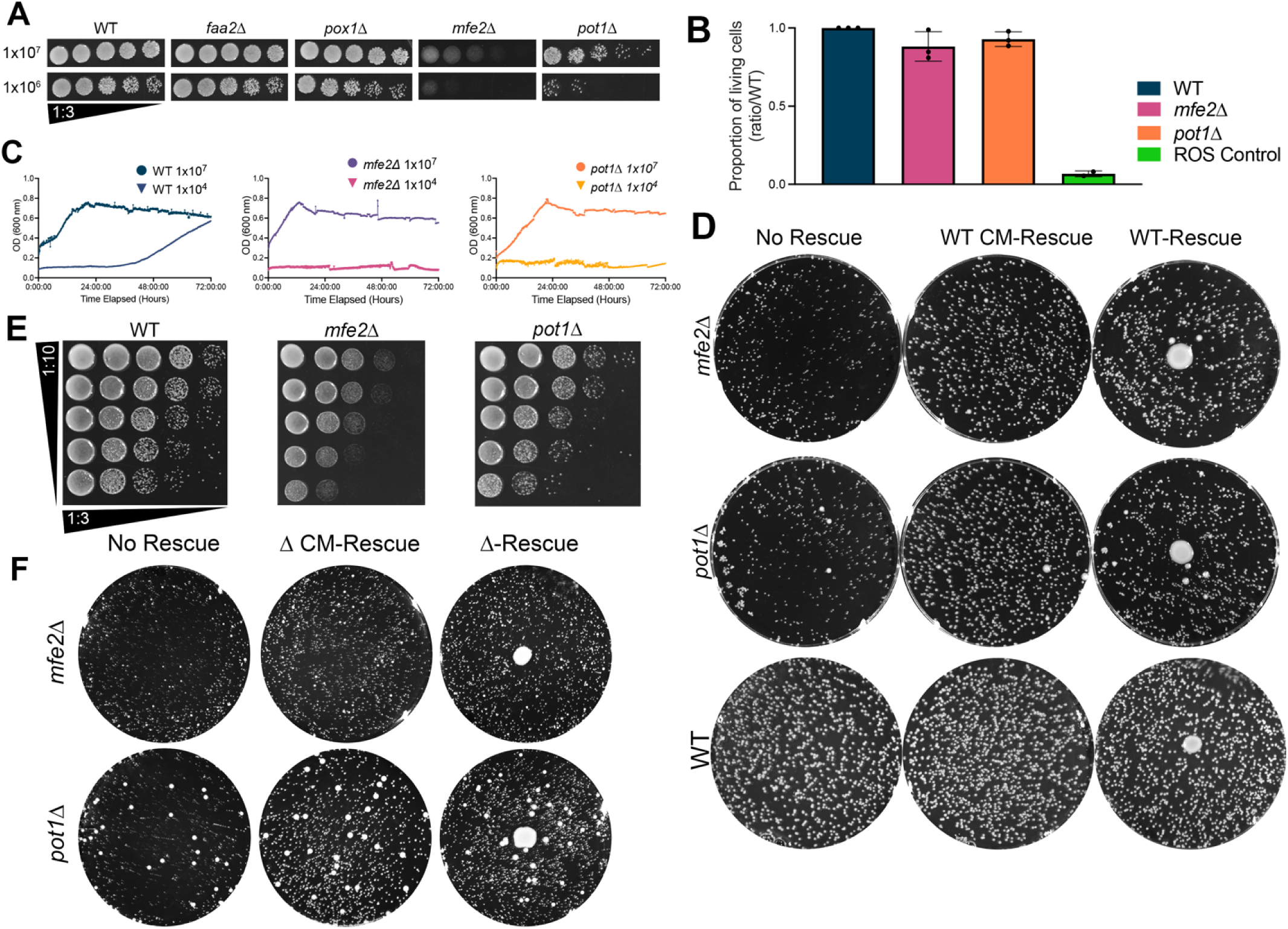
Cell density-related effects on growth. **(A)** Growth phenotypes were recapitulated when fungal cells were diluted from minimal media agar lawns. Mutants were precultured on minimal media agar lawns for 2-3 d, washed, counted, and diluted to two cell densities: 1x10^7^ and 1x10^6^ cells/mL. These two cell densities were serially diluted 1:3 onto minimal media plates for 48 h at 30°C. The growth defects of *mfe2*Δ and *pot1*Δ were partially rescued at 1x10^7^ cells/mL and exacerbated in the 1x10^6^ cells/mL serial dilution. **B)** Growth inhibition of *mfe2*Δ and *pot1*Δ did not correlate with a significant increase in cell death. Serial dilutions of 1x10^6^ cells/mL were spotted on minimal media agar supplemented with 2 μg/mL Phloxine and grown for 16 h at RT. Cell survival was measured using fluorescent cells as a readout for cell death and normalized to WT in each replicate (One-way ANOVA, ns). 5 mM H_2_O_2_-treated WT cultures were used as a positive control. **C)** Growth inhibition at low cell densities also occurs in liquid media. Growth curves (O.D. 600 nm) comparing cell densities 1x10^7^ cells/mL and 1x10^4^ cells/mL in minimal media over 72 h. **D)** WT-conditioned media and proximity to a WT colony rescued the growth of both mutants, indicating a secreted factor rescues growth. Left: 750 cells (75 μL of 1x10^4^ cells/mL) were spread on minimal media plates using glass beads. Middle: 1 mL of filtered WT conditioned media was applied to the plate. Right: approximately 1x10^6^ WT cells (10 μL of 1x10^8^ cells/mL) were spotted in the center of the agar dish. Plates were imaged after 4 d of growth at 30°C. **E)** *mfe2*Δ and *pot1*Δ show improved growth (relative to panel A) at lower cell densities when near higher cell density spots. We spotted 1:3 serial dilutions from cell densities ranging from 1x10^8^ to 1x10^4^ cells/mL on minimal media agar. Plates were imaged after 48 h at 30°C. **F)** *mfe2*Δ and *pot1*Δ produce the soluble factor necessary to rescue growth. Left: ∼750 cells (75 μL of 1 x 10^4^ cells/mL) were spread on minimal media agar plates. Middle: 1 mL of filtered mutant conditioned media was applied to the plate. Right: Mutants were self-rescued by spotting 10 μL of 1x10^8^ cells/mL (1x10^6^ total cells) mutant cells in the middle of the plate. Plates were imaged after 4 d at 30°C.

We considered that the reduction in growth was the result of a high rate of cell death (Fig. 4B). Phloxine B, a fluorescent yeast cell death indicator, was added to agar to measure cell death [40]. Serial dilutions of 1x10^6^ cells/mL were spotted on minimal media agar with 2 μg/mL Phloxine B. At least during short-term growth (16 h), increased cell death was not the explanation for the growth inhibition phenotype.

We examined growth in liquid cultures with two different starting inoculums: 1x10^7^ cells/mL and 1x10^4^ cells/mL (Fig. 4C). Growth inhibition at low cell densities also occurs in liquid media for *mfe2*Δ and *pot1*Δ. Additionally, growth curves performed at 1x10^6^ cells/mL and 1x10^5^ cells/mL showed intermediate, variable phenotypes, consistent with the variable virulence factor findings (Sup. Fig. 3).

Previous work showed that growth defects of *mfe2*Δ are rescued by supplementing mutants with nutrient-rich conditioned media made by WT cells or by spotting a WT rescue colony in the center of the agar plate [19]. However, it was not known if this phenotype would occur using minimal media and if these factors could rescue *pot1*Δ (Fig. 4D). We compared the growth of *mfe2*Δ and *pot1*Δ on untreated minimal media agar plates to both agar plates treated with 1 mL of filtered WT conditioned media and to agar plates with 1 x 10^6^ WT cells (10 μL of 1x10^8^ cells/mL) spotted in the center of the agar dish. WT-conditioned media and proximity to a WT colony rescued the growth defects of both mutants. Taken together, these results support the observation that peroxisomal β-oxidation mutants are rescued by a secreted factor [19].

Previous work also showed that in addition to long-range secretion, proximity to WT colonies rescued the growth of *mfe2*Δ during serial dilution experiments on YPD agar [19]. Our experiments suggested that the mutants can rescue themselves. To test if the mutants could also rescue themselves when grown near high cell density serial dilutions, we prepared cell suspensions ranging from 1x10^8^ cells/mL to 1x10^4^ cells/mL serial dilutions of all cell densities on the same agar dish (Fig. 4E). Both strains showed improved growth at lower cell densities when near higher cell density spots, indicating that *mfe2*Δ and *pot1*Δ could rescue themselves.

Next, we tested whether the mutant strains were capable of rescuing growth themselves via secretion from a mutant colony and mutant conditioned media (Fig. 4F). Such rescue would indicate that at high cell densities, these strains produce growth-rescuing metabolites. We repeated the experiments from panel D using conditioned media generated by the mutant strains and a mutant colony plated in the center. Both mutant conditioned media and mutant high-density colonies in the center of the plate were able to rescue the growth of the mutant strains. These data indicate that both *mfe2*Δ and *pot1*Δ produce a soluble factor that can rescue growth and thus are capable of “self-rescue”.

Curiously, we noted that colonies near the rescue colony were often smaller in size. This was counterintuitive as we expected that the colonies near the drop would be rescued more efficiently than the colonies in the periphery. We plated an increased volume of WT rescue colony (100 μL of 1x10^8^ cells/mL) (Sup Fig. 2). This produced a zone of mutant growth inhibition (red circle) surrounding the large WT rescue colony. This zone of inhibition was not present in the WT plate. To quantify the zone of inhibition, colony size measurements were compared between colonies close to the rescue colony and those on the outer zone (Sup Fig. 2). For both strains, the inner colonies were significantly smaller in size (Kruskall-Wallis test, ****, p-value<0.0001). There were no significant differences when comparing either the inner or outer zones of both strains to each other.

### Higher cell density rescued key cryptococcal virulence factors

We hypothesized that the growth defects of *mfe2*Δ and *pot1*Δ strains caused the variability of our virulence factor assays. Given the ability of both peroxisomal β-oxidation mutants to rescue their own growth defect, we wondered if we could restore virulence factor phenotypes by increasing the cell density. We first tested the ability of increasing cell density to rescue urease activity (Fig. 5A). We compared serial dilutions in the center of a Christensen’s urea agar plates and measured urease halo sizes (Fig. 5B). Comparing halo sizes produced by the lowest cell density tested, WT halos were significantly larger than the halos of both *mfe2*Δ (n=3 biological replicates, 2-way ANOVA, ****, p<0.0001) and *pot1*Δ (2-way ANOVA, **, p=0.0033). The halo sizes produced by the highest cell density tested did not differ between WT and *pot1*Δ and differed less significantly between WT and *mfe2*Δ (2-way ANOVA, *, p=0.0245).

**Figure 5.**
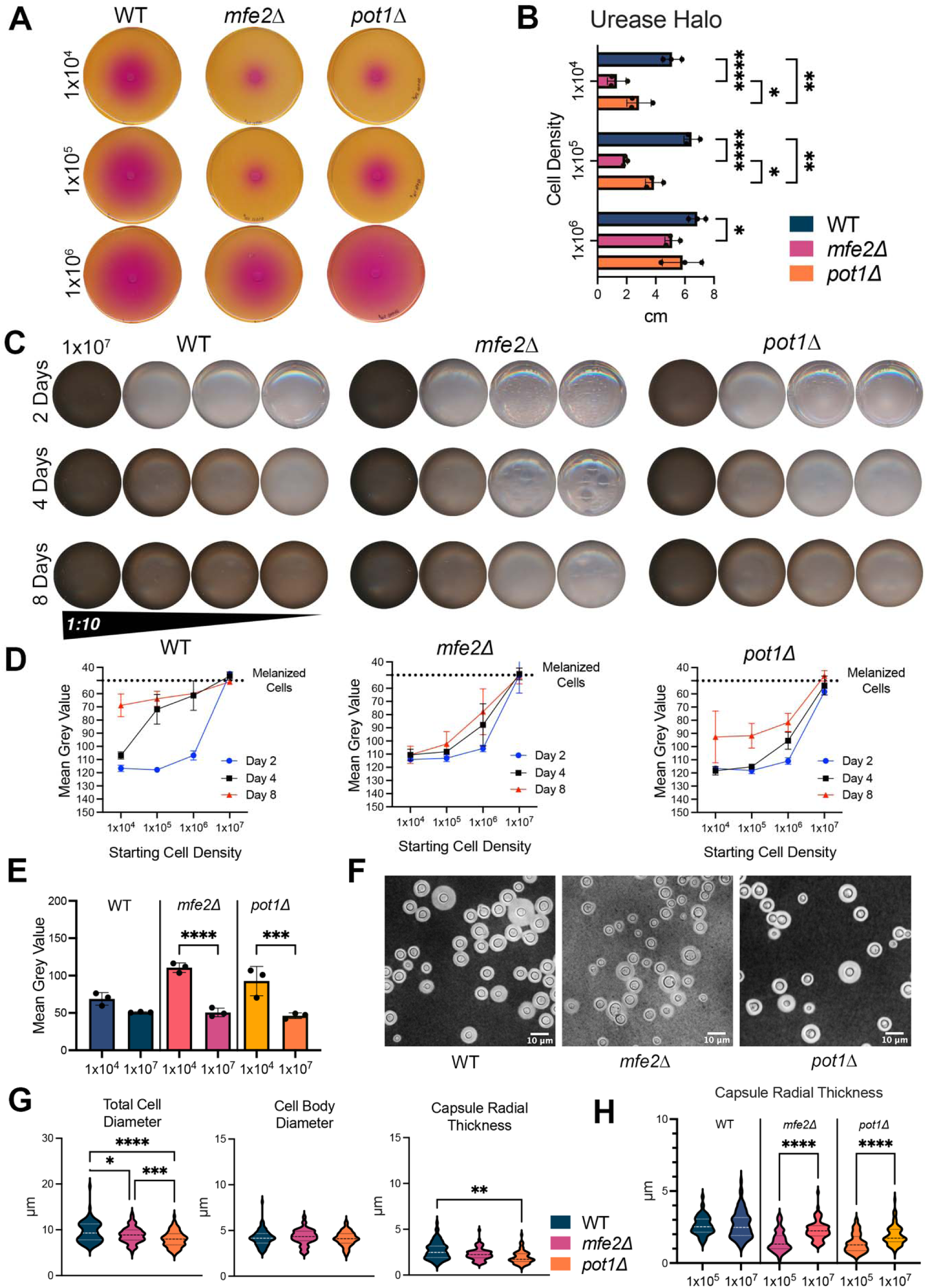
Higher cell density rescued key cryptococcal virulence factors. **(A)** Urease activity produced by various cell densities. A 10 μL volume with 1x10^6^, 1x10^5^, and 1x10^4^ cells in each spot were placed on the center of a Christensen’s urea agar plate for 48 h at 30°C. Representative images are shown. **B)** Urease halo size produced by various cell densities after 48 h (n=3 biological replicates). Comparing halo sizes from the 10 μL volume of 1x10^5^ cells/mL, WT halos were significantly larger than the halos of both *mfe2*Δ (2-way ANOVA, ****, p<0.0001) and *pot1*Δ (2-way ANOVA, **, p=0.0033). Comparing halos produced by the 10 μL volume of 1 x 10^8^ cells/mL, urease halo size did not differ between WT and *pot1*Δ and differed less significantly between WT and *mfe2*Δ (2-way ANOVA, *, p=0.0245). **C)** Melanization after 2, 4, and 8 d at various initial inoculua. Cryptococcal cultures were inoculated with cell densities of 1x10^7^, 1x10^6^, 1x10^5^, and 1x10^4^ cells/mL in minimal media supplemented with 1 mM L-DOPA. Representative images are shown. **D)** Mean grey value of melanization cultures over time from the various starting inoculua. *mfe2*Δ and *pot1*Δ showed delayed growth and melanization at lower cell densities. (n=3 biological replicates) **E)** Mean grey value comparisons at day 8. *mfe2*Δ and *pot1*Δ showed a significant reduction in melanization when the starting inoculum was 1x10^4^ cell/mL compared to 1x10^7^ cell/mL (n=3 biological replicates, one-way ANOVA, ****, p < 0.0001, ***, p = 0.0002). WT melanization was not significantly different. **F)** Representative counterstained India ink images from 1x10^7^ cell/mL starting inoculum after 7 d in minimal media without L-DOPA (40x magnification, scale bar 10 μm). **G)** Measurements of total cell diameter (capsule + cell body), cell body diameter, and capsule radial thickness (calculated from total cell diameter and cell body diameter). WT total cell diameter was significantly different from *pot1*Δ and *mfe2*Δ (n=3 biological replicates, ∼25 cells measured per replicate pooled, One-way ANOVA, *, p = 0.0105,****, p < 0.0001). Cell body sizes were not significantly different between the three strains. WT capsule sizes were not significantly larger than *mfe2*Δ but they were significantly larger than *pot1*Δ (One-way ANOVA, **, p = 0.0025). **H)** Comparing capsule radial thickness from Fig. 2F (2 x 10^4^ cells/mL) to capsule thickness from Fig. 4G (1 x 10^7^ cells/mL) revealed an increase in capsule thickness for both mutants at higher cell densities (One-way ANOVA, ****, p<0.0001). WT capsule radial thickness was not significantly different between the two experiments.

Next, we tested whether melanization could be rescued by increasing cell densities. We compared 1:10 serial dilutions in minimal media supplemented with 1 mM L-DOPA. Cell cultures were imaged after 2, 4, and 8 d (Fig. 5C), and mean grey values were measured (Fig. 5D). Strains *mfe2*Δ and *pot1*Δ showed delayed growth and melanization at lower cell densities. At day 8, strains *mfe2*Δ and *pot1*Δ showed a significant reduction in melanization (Fig. 5E) when the lowest starting inoculum was compared to the highest starting inoculum (n=3 biological replicates, one-way ANOVA, ****, p<0.0001, ***, p=0.0002). In contrast, WT melanization was not significantly different.

Additionally, increasing cell densities rescued capsule production (Fig. 5F). We measured total cell diameter (capsule and cell body), cell body diameter, and capsule radial thickness (calculated from total cell diameter and cell body diameter (Fig. 5G). WT total cell diameter was significantly different from *pot1*Δ and *mfe2*Δ (∼25 cells measured per biological triplicate, pooled, One-way ANOVA, *, p = 0.0105, ****, p < 0.0001). Cell body sizes were not significantly different between the three strains. WT capsule sizes were not significantly larger than *mfe2*Δ but they were significantly larger than *pot1*Δ (One-way ANOVA, **, p = 0.0025). Comparing capsule radial thickness from Fig. 2F to capsule thickness in Fig. 5G revealed an increase in capsule thickness for both mutants at higher cell densities (Fig. 5H) (One-way ANOVA, ****, p<0.0001). WT capsule radial thickness was not significantly different between the two experiments.

### RTG signaling controls the growth phenotypes of *pot1*Δ and *mfe2*Δ

In *S. cerevisiae,* RTG pathway mutants are unable to grow on acetate as a sole carbon source [41]. The *mfe2*Δ strain was also unable to utilize acetate as a sole carbon source [19]. We were curious if increasing the cell density would restore growth. To test the ability of mutants to grow on acetate as a sole carbon source with increased cell density, we substituted the 15 mM glucose for 15 mM sodium acetate and spotted 1:3 serial dilutions of two cell densities (Fig. 6A). At the higher cell density (left), no growth difference between the strains was observed, consistent with a cell-density secretion rescue. At the lower cell density (right), growth inhibition was exacerbated in both mutant strains relative to WT.

**Figure 6:**
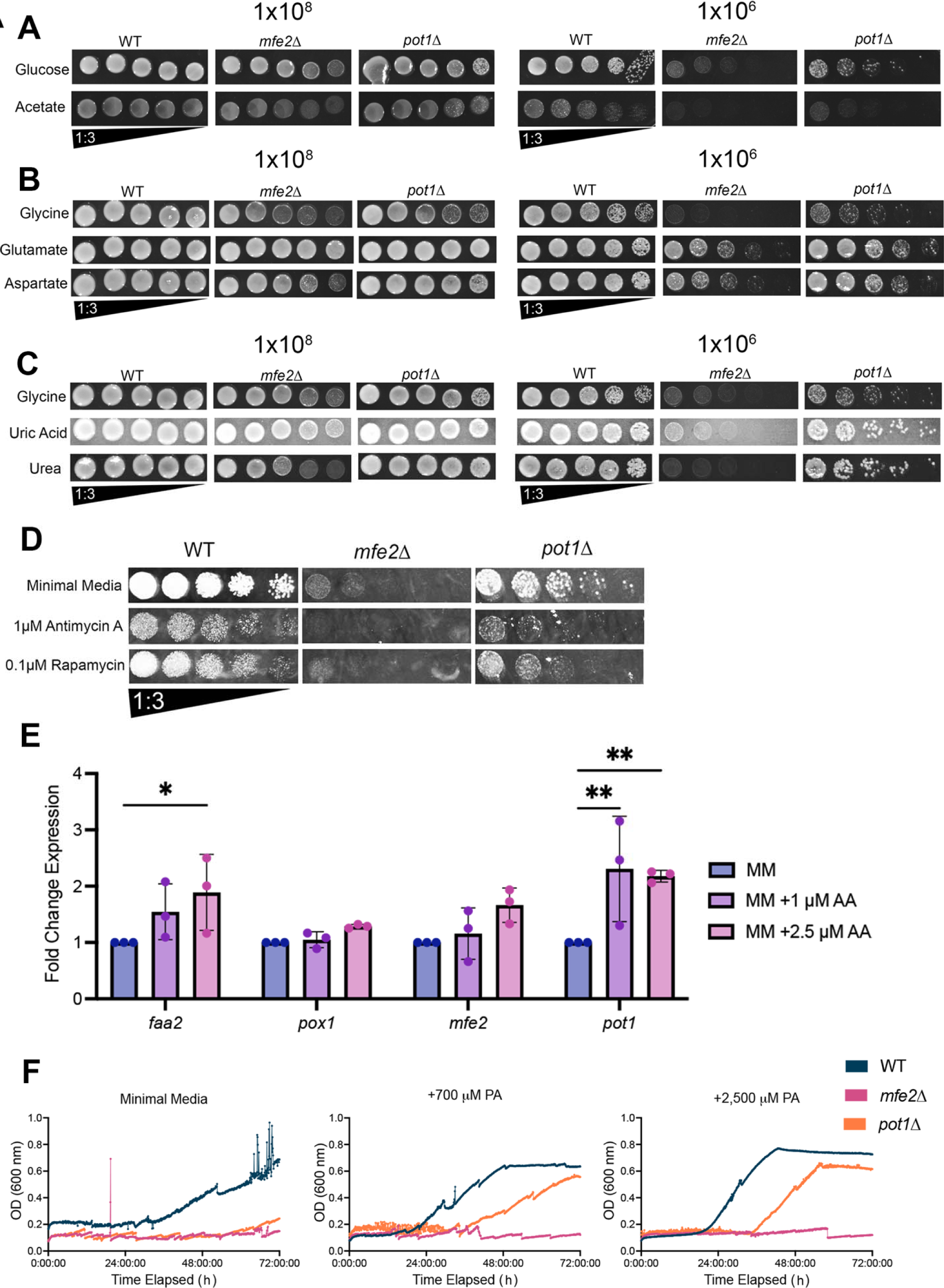
RTG signaling controls growth phenotypes of *pot1*Δ and *mfe2*Δ. **(A)** Substituting 15 mM glucose in minimal media with 15 mM sodium acetate inhibited the growth of both mutants at lower cell densities. (48 h of growth at 30°C, 15 mM sodium acetate). At 1x10^8^ cells/mL (left), no growth defects was observed. At 1x10^6^ cells/mL (right), growth inhibition was exacerbated in both mutant strains. **(B)** Substituting 13 mM glycine for 13 mM L-glutamic acid (glutamate) or L-aspartic acid (aspartate) rescued growth at low cell densities (48 h at 30°C). Similarly to acetate, at 1x10^8^ cells/mL (left) no growth defects was observed. At 1x10^6^ cells/mL (right), swapping glycine with glutamate or aspartate markedly rescued the growth of both mutant strains **(C)** Growth on environmental nitrogen sources. Substituting 13 mM glycine for 13 mM uric acid or urea did not rescue or exacerbate mutant growth defects. Two cell densities (1x10^8^ and 1x10^6^ cells/mL) were serially diluted 1:3 and spotted on agar for 48 h at 30°C. **(D)** Susceptibility to Antimycin-A and Rapamycin exposure. *mfe2*Δ and *pot1*Δ were susceptible to 1 μM antimycin-A and 0.1 μM Rapamycin exposure. **(E)** qPCR comparison of peroxisomal β-oxidation pathway gene expression after 2 h antimycin-A exposure in WT (KN99α). Fold-change expression relative to minimal media (blue). 1 μM antimycin-A (lilac), and 2.5 μM antimycin-A (pink). (n=3 biological replicates. 2-way ANOVA, Dunnett’s multiple comparisons test. *, p=0.0217, **, p=0.0010, **, p=0.0026). Antimycin-A exposure significantly induced expression of *pot1* at both concentrations tested. **(F)** Addition of pantothenic acid (PA) to the growth media rescues growth of *pot1*Δ but not *mfe2*Δ. Strains were seeded at 1x10^4^ cells/mL minimal media.

Glutamate and aspartate are RTG pathway inhibitors in *S. cerevisiae* [41]. Supplementation of either glutamate or aspartate restores the growth of RTG pathway mutants [41]. *C. neoformans* secretes metabolites involved in alanine, aspartate and glutamate metabolism during *in vitro* infection conditions [42]. We repeated the experiment in panel A, substituting the 13mM glycine for 13mM L-glutamic acid (glutamate) or L-aspartic acid (aspartate) (Fig. 5B). As expected, at the higher cell density no growth phenotype was observed. Interestingly, at the lower cell density, swapping glycine with glutamate or aspartate markedly rescued the growth of both mutant strains. The transcriptomic datasets and our growth phenotypes suggest that RTG signaling is repressed during log-phase growth. We hypothesized that the *pot1*Δ and *mfe2*Δ strains require secreted RTG-repressing metabolites to enter log-phase growth. Taken together, these experiments implicate retrograde signaling in the growth defects of peroxisomal β-oxidation genes.

To further validate the role of RTG signaling in the growth phenotypes, we tested other nitrogen source inducers of the RTG pathway. In *S. cerevisiae,* ammonia and urea activate RTG signaling [39]. *C. neoformans* secretes ammonia [43] and ammonia metabolism plays an important role in virulence [44, 45]. Furthermore, bird guano, a common *C. neoformans* environmental niche, is nitrogen-rich and has been reported to release ammonia [46]. We tested the ability of the environmentally relevant nitrogen sources urea and uric acid to induce the growth inhibition phenotypes of the *mfe2*Δ and *pot1*Δ. We substituted the 13 mM glycine in minimal media for 13 mM uric acid or urea (Fig. 5C). We found that, in addition to glycine, *mfe2*Δ and *pot1*Δ have growth defects on uric acid and urea at low cell densities.

We also tested drugs known to induce RTG signaling. Antimycin-A inhibits the mitochondrial electron transport chain. If RTG signaling was activated in the *pot1*Δ and *mfe2*Δ strains, antimycin A would be expected to induce growth defects. We found that 1 μM antimycin-A inhibited the growth of *pot1*Δ and *mfe2*Δ (Fig. 5D). In *S. cerevisiae,* Rapamycin activates the RTG pathway [47, 48]. We found that exposure to 0.1 μM Rapamycin inhibited growth of *pot1*Δ and *mfe2*Δ.

We previously showed that nickel induced the expression of the peroxisomal β-oxidation pathway. If RTG signaling underlies these phenotypes, then exposure to Antimycin-A should also induce gene expression. Using quantitative PCR, we compared the expression of the peroxisomal β-oxidation pathway genes in WT cultures after 2 h of 1 μM or 2.5 μM Antimycin-A exposure (Fig. 5E). We found that expression of *pot1* increased significantly in a dose-dependent manner.

In *C. neoformans,* pantothenic acid (Vitamin B5), the precursor for Coenzyme A (CoA), plays a role in quorum sensing and rescues fungal cells from a viable-not-culturable phenotype [24, 42, 49, 50]. Given the fact that RTG signaling is inhibited by a functioning TCA cycle, we wondered if pantothenic acid supplementation could rescue the growth of *pot1*Δ and *mfe2*Δ. Addition of pantothenic acid rescued the growth of *pot1*Δ, but not *mfe2*Δ (Fig. 5F). Increasing the concentration of pantothenic acid did not rescue the growth of *mfe2*Δ. Taken together, these findings show RTG signaling controls the growth defects of *pot1*Δ and *mfe2*Δ and implicate RTG and peroxisomes as required for *C. neoformans* growth in multiple environments. Furthermore, we show that growth inhibition is rescued by increasing cell density, further illustrating the importance of a secreted factor for the *pot1*Δ and *mfe2*Δ strains to enter log-phase.

### *mfe2*Δ and *pot1*Δ maintain large peroxisomes over time

In several yeast species, Pot1 and Mfe2 (also Fox2) regulate peroxisome size and abundance [30, 51, 52]. We hypothesized that enlarged peroxisomes might underly the growth defects. To test this, we compared peroxisome induction between 2 and 16h after growth in minimal media using transmission electron microscopy (TEM) (Fig. 7A).

**Figure. 7.**
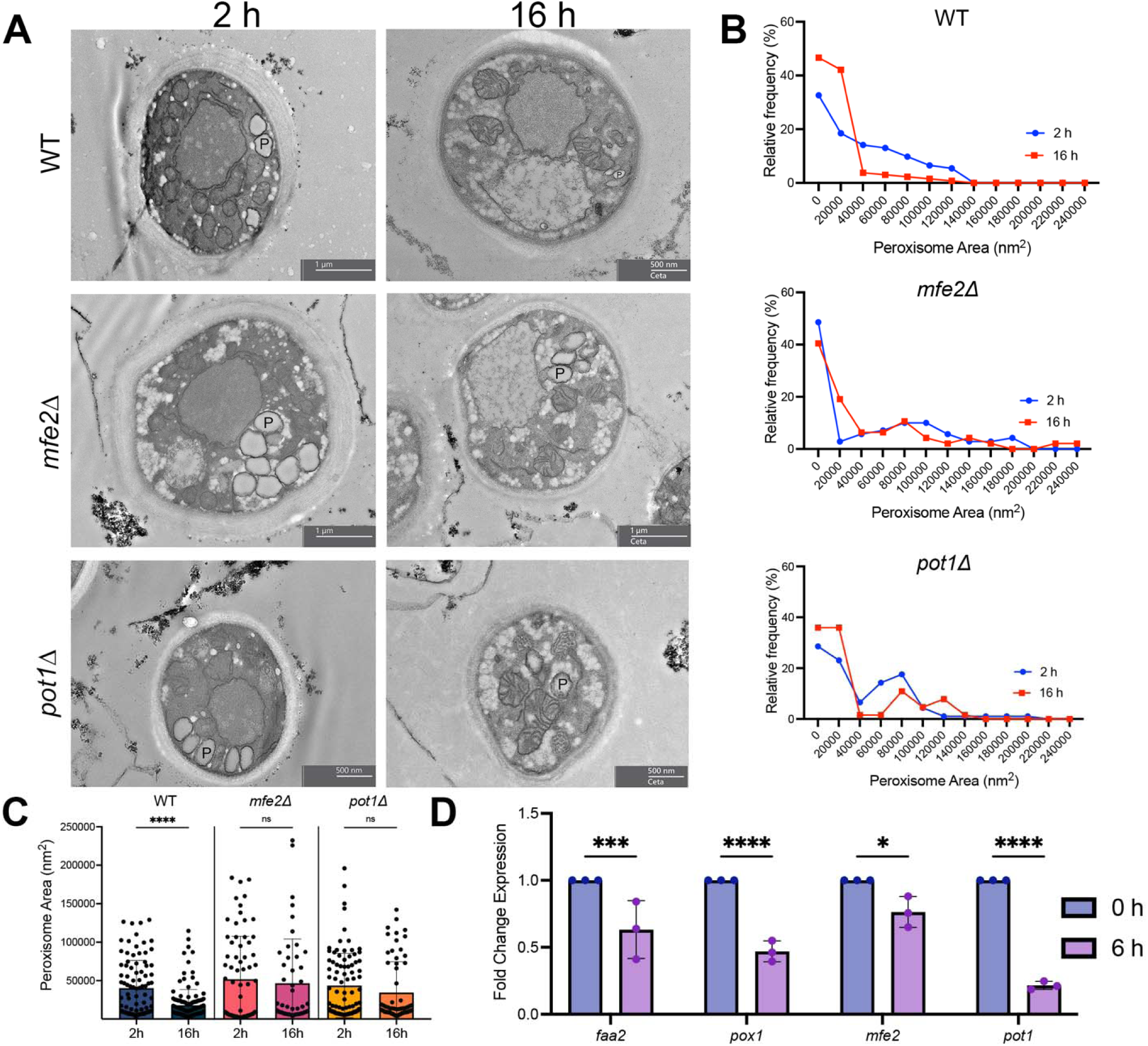
*mfe2*Δ, and *pot1*Δ maintain large peroxisomes over time.. **A)** Representative TEM images of WT, *mfe2*Δ, and *pot1*Δ after 2 h and 16 h of growth in minimal media. P indicates peroxisome. **(B)** Relative frequency of peroxisomal areas at 2 h (blue) and 16 h (red). **(C)** Comparison of peroxisome areas between 2 h and 16 h. WT peroxisomal area significantly decreases over time (Kruskal-Wallis p < 0.0001). **(D)** qPCR comparison of peroxisomal β-oxidation pathway gene expression between 0 h and 6 h of growth in minimal media. (n=3 biological replicates. 2-way ANOVA, Sidak’s multiple comparisons test, *, p= 0.0228, ***, p=0.0006, ****, p<0.0001).

At 2h, growth in minimal media induced peroxisomes of varying sizes across all strains, including WT (Fig. 7A). At 16 h, the peroxisomal area of *mfe2*Δ, and *pot1*Δ was increased relative to WT (Fig. 7 B and C). We hypothesize that while WT, *mfe2*Δ, and *pot1*Δ strains manifested large peroxisomes initially, the *mfe2*Δ and *pot1*Δ strains may be unable to exit this growth state, accumulating toxic intermediates and enlarged peroxisomes over time. Additionally, we compared the following 3 carbon and nitrogen source combinations at 2 h: glucose/glycine, acetate/glycine, glucose/glutamate (Sup Fig. 3). We found that across carbon and nitrogen sources tested all strains induced peroxisomes at 2 h.

To confirm that expression of the β-oxidation pathway genes is repressed during log-phase growth, we used qPCR to compare relative expression at 0 and 6h of growth in minimal media. We found that expression of β-oxidation pathway genes decreased significantly between 0 h and 6 h of growth in minimal media (Fig. 7D). These results are in agreement with the previously observed expression patterns of *MFE2* expression in YPD over time [19].

### *mfe2*Δ and *pot1*Δ had attenuated virulence in the mouse model

First, we performed *Galleria mellonella* infections to assess the virulence of *mfe2*Δ and *pot1*Δ. Compared to WT, only *mfe2*Δ was significantly avirulent in this model system (Log-rank test: WT vs. *mfe2*Δ*, ***,* P value = 0.0004).

Previously, the cryptococcal *mfe2*Δ strain was shown to have attenuated virulence using the mouse inhalation model of infection [19]. Here, we tested the virulence of *mfe2*Δ and *pot1*Δ using the intravenous model of infection (Fig. 8B). We found that both *mfe2*Δ and *pot1*Δ had significantly delayed virulence compared to WT (Log-rank test: WT vs. *mfe2*Δ*, ***,* P value = 0.0004, WT vs. *pot1*Δ*, ***,* P value = 0.0008).

**Figure 8.**
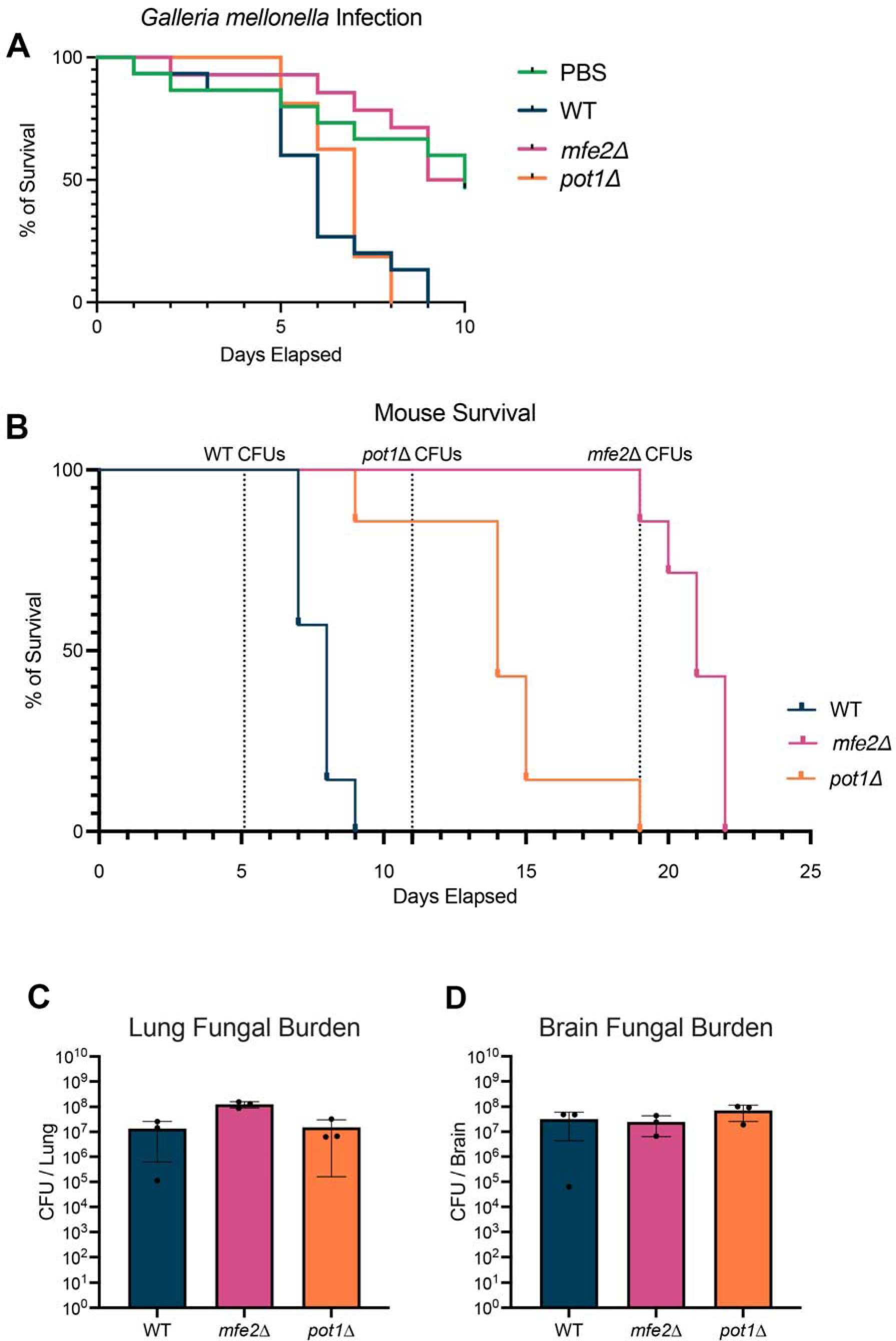
*mfe2*Δ and *pot1*Δ have delayed virulence. **(A)** *Galleria mellonella* survival. n = 15 larvae per fungal strain. (Log-rank test: WT vs. *mfe2*Δ*, ***,* adj P value = 0.0004) **(B)** Mouse survival post IV infection. *mfe2*Δ and *pot1*Δ had significantly attenuated virulence compared to WT (n = 7 mice per fungal strain. WT vs. *mfe2*Δ Log-rank adj P value = 0.0004, WT vs. *pot1*Δ Log-rank adj P value = 0.0008). CFUs were collected for each strain at the days indicated with dotted lines. (WT on day 5, *mfe2*Δ on day 19, *pot1*Δ on day 11) **(C&D)** CFUs post IV infection. CFUs were obtained as indicated in panel C (WT on day 5, *mfe2*Δ on day 19, and *pot1*Δ on day 11). Kruskal-Wallis test found no significant differences. Recovered CFUs were plated on glutamate-substituted minimal media plates and incubated at 30°C for 2 d for WT and *pot1*Δ and 4 d for *mfe2*Δ.

Previously, there was a significant reduction in *mfe2*Δ brain CFUs post-intranasal infection [19]. It was unknown whether this phenotype was the result of an inability to colonize the brain tissue or to disseminate. Our results suggest that this phenotype could be explained by delayed growth *in vivo* or by the previous work plating of CFUs on YPD plates lacking RTG repressing metabolites. We collected brain and lung CFUs at comparable time points during intravenous infection progression to assess the ability of the *mfe2*Δ and *pot1*Δ strains to colonize these tissues (Fig. 8 C and D). Recovered CFUs were plated on glutamate-substituted minimal media plates, as these conditions rescued the growth of both strains. By collecting CFUs on the same day before most deaths occurred, this allowed us to compare a similar time point in infection of each strain. CFUs were obtained for WT on day 5, *pot1*Δ on day 11, and *mfe2*Δ on day 19. We found no significant differences between CFU burdens in the brain or lung, indicating that both peroxisomal mutants can colonize these tissues, albeit at a slower pace than WT. Our work shows that peroxisomal function and RTG contribute to optimal growth within mammalian tissues and are important components of *C. neoformans* metabolic flexibility.

## Discussion

Functioning mitochondria are essential in cryptococcal virulence, stress tolerance, and antifungal susceptibility [15, 53]. By exposing *C. neoformans* cultures to a low concentration of nickel, a known hypoxia-mimetic and mitochondrial inhibitor in yeast [20–23, 53, 54], we found that nickel exposure upregulated genes including peroxisomal fatty acid β-oxidation pathway, peroxisome biogenesis, and amino acid metabolism. Here, we studied this previously unexplored connection between cryptococcal respiration inhibition and the peroxisomal fatty acid β-oxidation. In summary, we show that RTG signaling plays a novel role in cell density-dependent metabolic reprogramming of *C. neoformans*.

In *Saccharomyces cerevisiae*, mitochondrial retrograde signaling (RTG) is a pro-survival pathway where mitochondrial dysfunction, among other stressors, signals to the nucleus to induce the expression of genes in the RTG pathway [55]. RTG signaling is induced by acetate, urea, mitochondrial inhibition, and rapamycin [47, 56–58]. In agreement with this literature, we found that cell-density-dependent growth defects occurred in both *mfe2*Δ and *pot1*Δ in the presence of acetate, urea, mitochondrial inhibition, and rapamycin. Conversely, *S. cerevisiae* RTG signaling is repressed by nitrogen sources glutamate and aspartate, which are anaplerotic molecules. We found that growth was rescued in both *mfe2*Δ and *pot1*Δ when the strains were exposed to these known RTG repressors. These results support that the RTG pathway controls the growth phenotypes in both *mfe2*Δ and *pot1*Δ.

Retrograde signaling in *S. cerevisiae* is controlled by two helix-loop-helix transcription factors (*RTG1* and *RTG3*) and an ATP-binding regulatory protein (*RTG2*) [34, 41]. *C. neoformans* lacks known homologs of *RTG1* and *RTG2* [36], but does possess an *RTG3* homolog known as *MLN1* [36, 37]. Prior to this work, the existence of RTG signaling had not been characterized in *C. neoformans*, in part due to the lack of sequence homology in the regulatory factors *RTG1, RTG2,* and *RTG3.* Future work will characterize the regulatory factors in the *C. neoformans* RTG signaling pathway.

The basic premise of mitochondrial retrograde signaling is conserved in other eukaryotes, including humans [59]. However, the specific metabolic signals and upstream signaling pathways are not well conserved between organisms [35, 59]. As evidence of this, in the pathogenic yeast *Candida albicans* the RTG pathway is regulated by TOR, carbon source, and sphingolipid content [60, 61]. Hence, the regulation and function of this pathway in *C. neoformans* may also be unique.

Importantly, we discovered that not only was the growth inhibition phenotype of *mfe2*Δ and *pot1*Δ cell-density-dependent, but also that *mfe2*Δ and *pot1*Δ produce secreted metabolites necessary for rescuing growth and/or breaking down inhibitory factors. Hence, by increasing the cell density of *mfe2*Δ and *pot1*Δ, we were able to use the “self-rescue” phenotype to fully or partially rescue all virulence factor defects and reduce variability. Interestingly, the cryptococcal *mfe2*Δ strain was also rescued by growing near an *E. coli* colony [19]. As such, the secreted product in this system is unlikely to be a *C. neoformans*-specific molecule.

Our data indicates that peroxisomal β-oxidation is induced during low-cell-density conditions and that growth is rescued by secreted RTG repressing metabolites. In *C. neoformans,* metabolite secretion is influenced by both cell density and time [42]. Exposure to these metabolites would allow fungal cells to enter log-phase where RTG signaling appears to be repressed. During *in vitro* infection conditions, *C. neoformans* secretes RTG repressing metabolites involved in alanine, aspartate, and glutamate metabolism [42]. Interestingly, at higher cell-density (MOI 100 vs. MOI 10), the abundance of RTG repressing metabolites are rapidly depleted over time and the presence of metabolites involved in β-Alanine metabolism increases [42]. Future work will definitively characterize the secretion of RTG repressing metabolites in this system.

The *C. neoformans* quorum sensing system is integrated with nutrient sensing [12], similar to several bacterial systems [62]. Our growth phenotypes mostly closely mirror those of the serotype D *tup1*Δ strain, where low cell density cultures are unable to grow and require supplementation of quorum-sensing peptide Qsp1 [63]. However, Qsp1, an important quorum sensing peptide in *C. neoformans* [13], is not responsible for the growth rescue of *mfe2*Δ in YPD-conditioned media. Our data suggests that amino acid availability plays an important role in the cell-density sensing of *C. neoformans* upstream of the RTG pathway. In *S. cerevisiae,* RTG-deficient strains exhibit significantly altered amino acid metabolism [64]. In agreement with this, we found that gene expression involved in the metabolism of amino acids (tryptophan phenylalanine and β-alanine) were increased during nickel exposure. This is consistent with *S. cerevisiae* metabolome analysis where the presence of alanine and the aromatics phenylalanine and tyrosine consistently increased with hypoxia [22]. We hypothesize that the “zone of inhibition” phenotype that occurs may be the result of the depletion of RTG repressing nitrogen sources and/or increased exposure to secreted β-Alanine from the rescue colony. Future work will explore the role of amino acid secretion in cell-density dependent sensing.

At high cell densities, *C. neoformans* secretes Pantothenic acid (vitamin B5), a precursor of Coenzyme A (CoA), early in infection conditions [42]. We found that the addition of Pantothenic acid rescued the growth of *pot1*Δ, but not *mfe2*Δ. Interestingly, biotin (vitamin B8/B7) partially rescued the growth defect of cryptococcal *mfe2*Δ [19]. Biotin is a coenzyme for several mitochondrial carboxylases with varying functions, including resupplying TCA cycle intermediates [65]. B vitamins are important regulators of mitochondrial-nuclear communication [65], and cellular CoA levels regulate the balance between carbohydrate and lipid metabolism [66]. We hypothesize that both biotin and Pantothenic acid supplementation support mitochondrial function and suppress the induction of RTG anaplerotic metabolic pathways but are not sufficient to fully rescue *mfe2*Δ.

Our data suggests that large peroxisomes cause the growth defects of *mfe2*Δ, and *pot1*Δ. In several yeast species, Pot1 and Mfe2 (Fox2 in *C. albicans*) regulate peroxisome size and abundance [30, 51, 52]. However, we did not observe growth inhibition in the *faa2*Δ or *pox1*Δ strains. Faa2 and Pox1 have not been implicated in peroxisome size or growth inhibition phenotypes in other yeasts [28, 29]. Interestingly, we found that peroxisomes were induced in WT, *mfe2*Δ, and *pot1*Δ after 2 hours in all minimal media tested (glucose/glycine, acetate/glycine, glucose/glutamate). While WT, *mfe2*Δ, and *pot1*Δ strains induced large peroxisomes initially, the later time point revealed that WT peroxisomes became smaller in size. We hypothesize that *mfe2*Δ and *pot1*Δ strains are unable to exit the initial metabolic state, likely accumulating toxic intermediates and enlarged peroxisomes over time. In other yeast species, the role of Pot1 and Mfe2 (alias Fox2) in peroxisome size and abundance regulation were assessed after overnight incubations [30, 51, 52]. Therefore, early peroxisomal induction in the WT strain would be missed. We hypothesize that the initial induction of peroxisomes is not dependent on functional Mfe2 or Pot1, but another upstream signal, such as RTG signaling. Interestingly, these same organelles were recently observed in *C. neoformans* cells, labeled as young vacuoles [67]. This work hypothesized that these small young vacuoles fuse to make the larger vacuoles observed in older cells. We posit that perhaps this organelle serves a metabolic function in the *C. neoformans* cell related to lipid and glucose metabolism.

In *C. albicans,* the RTG pathway is thought to have diverged as RTG mutants are not glutamate auxotrophs [68]. However, our work implies that RTG phenotypes may only be present in a cell density dependent manner. In support of this, *C. albicans mfe2*Δ (alias *fox2*Δ) and *pot*Δ*1* strains grown on acetate appear to show the same cell-density-dependent inhibition that we observe [69, 70]. Conversely, a glucose growth defect was not observed in these *C. albicans* strains [70], in contrast with our data, and supporting the hypothesis that the cryptococcal peroxisome plays a diverged role in RTG and glucose metabolism compared to *C albicans*.

We observed virulence attenuation of the *mfe2*Δ and *pot1*Δ strains. There are several, non-mutually exclusive, explanations for this: 1) dysfunctional and enlarged fungal peroxisomes prevented optimal growth within the host, 2) functional RTG pathways are necessary for early colonization and optimal growth in murine host, and 3) loss of RTG and peroxisomal function at low cell densities inhibited virulence factors necessary to evade the immune response, and delayed growth in the host. Phagocytosis induces immediate metabolic reprogramming in cryptococcal cells, including hypothesized RTG response pathways: glyoxylate pathway, gluconeogenesis, lipid catabolism and two-carbon metabolism [5–9]. Hence, RTG controlled metabolic pathways may be important for optimal growth within the host, but are partially compensated by other pathways, either by de novo synthesis or by sequestering of host compounds which feed compensatory pathways.

In summary, our work shows that *C. neoformans* has a previously uncharacterized RTG signaling response induced in low cell-density conditions. In the future, characterization of the retrograde signaling pathway regulators, potential diverged functions, and signaling metabolites are necessary to establish the importance of this response. The growth defects of *mfe2*Δ and *pot*Δ*1* that occur when RTG signaling is active make these strains promising candidates for exploring the induction and repression of RTG signaling in this fungus.

## Materials and Methods

### Ethical statement

All animal procedures were performed with prior approval from the Animal Care and Use Committee (ACUC) of Johns Hopkins University, under approved protocol number MO24H60. Handling of mice and euthanasia with CO_2_ in an appropriate chamber were conducted in accordance with guidelines on Euthanasia of the American Veterinary Medical Association. Johns Hopkins University is accredited by AAALAC International, in compliance with Animal Welfare Act regulations and Public Health Service (PHS) Policy and has a PHS Approved Animal Welfare Assurance with the NIH Office of Laboratory Animal Welfare.

### Fungal strains

Culture stocks were stored in 15% glycerol in YPD at 80°C. Wild-type *Cryptococcus neoformans* serotype A strain KN99α, *mfe2*Δ, and *pot1*Δ were obtained from the 2015 Fungal Genetics Stock Center gene deletion library generated by the Madhani Lab. All 2015 library collection strains were validated for NAT cassette insertion with primers corresponding to the cassette insertion location.

### Nickel exposure mRNA-Seq

*C. neoformans* cells were cultured in minimal media with a sub-inhibitory concentration of nickel sulfate (31.5 μM NiSO_4_) and was performed gene expression analysis using bulk RNA-Seq. *C. neoformans* was harvested from cultures. Pellets were washed with PBS, flash frozen, and stored at -80 °C till processing. Homogenization was performed in Trizol Reagent (ThermoFisher) using silica spheres (Lysing Matrix C, MP Biomedical) with shaking in a FastPrep 120 (MP BioMedical), at speed 6 for 30 sec, 4 times. Homogenates were held on ice between each cycle. Subsequent extraction and purification of total RNA was performed with the PureLink RNA Mini Kit (ThermoFisher), including the on-column PureLink DNAse treatment, according to manufacturer’s recommended protocol. Quantitation of RNA was performed using a NanoDrop 1000. Quality assessment was performed by RNA Nano LabChip Analysis in a BioAnalyzer 2100 (Agilent). Libraries for RNA-seq were prepared from 500 ng total RNA using the Illumina TruSeq RNA Library Prep kit v2, according to manufacturer’s Low Sample protocol (Illumina). Quality assessment of libraries was performed by High Sensitivity ScreenTape analysis on a TapeStation 2200 (Agilent). Quantitation of libraries was performed by PicoGreen Fluorescent detection in a Spectramax M2 plate reader. Libraries were diluted, pooled, and subsequently transferred to the JHMI High-Throughput Sequencing Core for paired end, 100 bp reads on an Illumina HiSeq 2500. Analysis was performed using Partek Flow NGS software, with RNA-seq Toolkit (Partek Inc). Alignment was performed with the STAR aligner to both the genome/transcriptome created from [71]. The analysis workflow included QC metrics, ‘Normalize counts’, GSA for differential read counts, and filtering to Gene Lists. Genes with 2-fold expression difference with p-value <0.05 were considered differentially regulated. Venn diagrams were used for the comparison of gene lists. Gene ontology analysis was performed by uploading the differentially regulated gene list into FungiDB with the GO in-built analysis.

### Media

For most experiments, minimal media was used (15.0 mM glucose, 10.0 mM MgSO_4_, 29.4 mM KH_2_PO_4_, 13.0 mM glycine, 3.0 mM vitamin B_1_, pH 5.5). For alternate carbon and nitrogen sources, the following substitutions were made: Acetate substituted minimal media (15.0 mM acetate, 10.0 mM MgSO_4_, 29.4 mM KH_2_PO_4_, 13.0 mM glycine, 3.0 mM vitamin B_1_, pH 5.5). Glutamate-substituted minimal media (15.0 mM glucose, 10.0 mM MgSO_4_, 29.4 mM KH_2_PO_4_, 13.0 mM glutamate, 3.0 mM vitamin B_1_, pH 5.5)). Aspartate-substituted minimal media (15.0 mM glucose, 10.0 mM MgSO_4_, 29.4 mM KH_2_PO_4_, 13.0 mM aspartate, 3.0 mM vitamin B_1_, pH 5.5). For virulence experiments, strains were pre-cultured from frozen stocks onto yeast extract-peptone-dextrose (YPD) medium.

### Growth conditions for screening experiments

For initial virulence factor and stress response screening, yeast strains were resuspended from minimal media agar in 96-well plates containing 200 μL minimal media at a cell density of 1x10^7^ cells/mL for 18 h. 10 μL of 1:3 serial dilutions were spotted directly onto minimal media agar. For endogenous ROS stress, 97% diethyl maleate (DEM) (Millipore Sigma) was added directly to cooled agar, and plates were used immediately. Fluconazole (FLC) (MilliporeSigma) was added directly to cooled agar, and plates were used immediately. FLC stock solutions were maintained in DMSO at a concentration of 5mg/mL. To test the impact of low cell density on growth, the cell density was reduced to 1x10^4^ cells/mL in 96-well plates containing 200 μL minimal media per well. 10 μL of 1:3 serial dilutions were spotted directly onto minimal media agar. Plates were imaged after 48 h of growth at 30°C.

### Growth conditions for virulence and growth rescue experiments

Frozen stocks were streaked directly onto minimal media agar at lawn density and incubated at 30°C for 2-3 d. To assay growth differences on agar, fungal cells in the minimal media agar lawns were resuspended in PBS, washed, counted, and diluted to two cell densities: 1x10^7^ cells/mL and 1x10^6^ cells/mL. These two cell densities were serially diluted 1:3 and spotted onto minimal media plates for 48 h at 30°C. To assay growth differences in liquid media, 1x10^7^ cells/mL and 1x10^4^ cells/mL were seeded into 12-well dishes containing 1 mL of minimal media each. O.D. 600 was measured every 5 minutes using the SpectraMax M5 plate reader (Molecular Devices) at a temperature of 30°C with shaking between reads.

### Urease secretion assay

Initial urease secretion experiments were performed by spotting 10 μL of cell suspension from 3x10^6^ cells/mL (3x10^5^ total cells) on Christensen’s urea agar for 48 h at 30°C. Plates were scanned using a CanoScan9000F scanner at 600 dpi. Urease halos were measured using the Measure tool on FIJI image processing software. Cell density urease secretion experiments were performed by spotting 10 μL of 1x10^6^, 1x10^5^, and 1x10^4^ cells on Christensen’s urea agar for 48 h at 30°C. Plates were scanned using a CanoScan9000F scanner at 600 dpi. Urease halos were measured using the Measure tool on FIJI image processing software.

### Melanization assay

Initial melanization experiments were performed with cultures grown for 7 d in 5 mL of minimal media (2x10^4^ cells/mL) supplemented with 1 mM L-3,4-dihydroxyphenylalanine (L-DOPA). Cells were grown under continuous culture wheel rotation at 30°C. After 7 d, 2 mL of each culture were placed in a 12-well plate and scanned with a CanoScan9000F scanner at 600 dpi. Mean Gray Value for each well of culture was determined using the Measure tool on FIJI image processing software [36]. Cell density melanization experiments were performed by inoculating cell densities of 1x10^7^, 1x10^6^, 1x10^5^, and 1x10^4^ cells/mL in 12-well plates containing 2 mL minimal media supplemented with 1 mM L-DOPA. Cell cultures were grown under continuous culture wheel rotation at 30°C. Cell cultures were imaged after 2, 4, and 8 d with a CanoScan9000F scanner at 600 dpi. Mean Gray Value for each well of culture was determined using the Measure tool on FIJI image processing software [36].

### Capsule growth assay

Initial capsule experiments were performed with cultures grown for 7 d in 5 mL minimal media (2 x 10^4^ cells/mL) without L-DOPA. Cells were grown under continuous culture wheel rotation at 30°C. Capsule was visualized with India ink negative staining after 7 d of growth using 5 μL of cell suspension and 2 μL of India ink. Images were obtained using an Olympus AX70 microscope (Olympus, Center Valley, PA) using a 40X objective. Total cell diameter (capsule and cell body), cell body diameter, and capsule radial thickness (calculated from total cell diameter and cell body diameter) were determined using the Measure tool on FIJI image processing software. Cell density capsule growth experiments were performed by inoculating cell densities of 1x10^7^ cells/ mL in 5 mL minimal media. Cells were grown under continuous culture wheel rotation at 30°C. Capsule was visualized with India ink negative staining after 7 d of growth using 5 μL of cell suspension and 2 μL of India ink. Images were obtained using an Olympus AX70 microscope (Olympus, Center Valley, PA) using a 40X objective and measured in FIJI. We measured total cell diameter (capsule and cell body), cell body diameter, and capsule radial thickness (calculated from total cell diameter and cell body diameter.

### Phloxine B staining

Serial dilutions of 1x10^6^ cells/mL were spotted on minimal media agar containing 2 μg/mL Phloxine B (85%, ThermoFisher). Plates were used directly after Phloxine B addition and kept shielded from light to avoid photo bleaching. After 16 h of growth at room temperature, cell survival was assessed using fluorescent cells as a readout for cell death and normalized to WT in each replicate. Images were obtained using the Leica Thunder Imaging system directly imaging the colonies on Phloxine B containing agar as first described in [72]. ROS treated WT cells were used as a positive control.

### Conditioned media and long-range secretion rescue

Conditioned media (CM) was prepared from 5 mL minimal media cultures seeded at a starting density of 1x10^7^ cells/mL. After 5 days of growth at 30°C, cells were spun down and the supernatant was filtered using a 0.2 µm filter. CM was used for experiments immediately. For untreated plates, a total of 750 cells (75 μL of 1x10^4^ cells/mL) were spread on minimal media plates using glass beads. For conditioned media rescue, 1 mL of filtered WT, *pot1*Δ, or *mfe2*Δ conditioned media, prepared as stated above, was applied uniformly to the plate surface before cells were added using glass beads.

To assess the ability of a high-density colony in the center of the plate to growth, 1x10^6^ WT or mutant cells (10 μL of 1x10^8^ cells/mL) were spotted in the center of the agar dish. Plates were imaged after 4 d of growth at 30°C.

To test if the mutants could also rescue themselves when grown near high cell density serial dilutions, we prepared cell suspensions ranging from 1x10^8^ cells/mL to 1x10^4^ cells/mL (1:10 dilutions) and spotted 1:3 serial dilutions of all cell densities on the same agar dish. Plates were imaged after 48 h on minimal media agar.

### Zone of inhibition

To assess the ability of a larger high-density colony in the center of the plate to inhibit growth, 100 μL of 1x10^8^ cells/mL WT or mutant cells were spotted in the center of the agar dish. Plates were imaged after 4 d of growth at 30°C. To quantify the zone of inhibition, colony size measurements were compared between colonies close to the rescue colony and those on the outer zone using FIJI.

### Pantothenic acid supplementation

To assay growth differences in the presence of pantothenic acid, 1x10^4^ cells/mL were seeded into 12-well dishes containing 1 mL of minimal media each. Wells were supplemented with 700 or 2,500 μM Pantothenic acid (PA). OD 600 was measured every 5 minutes using the SpectraMax M5 plate reader (Molecular Devices) at a temperature of 30°C with shaking between reads.

### Rapamycin and Antimycin-A treatment

Rapamycin stocks were maintained at -20°C at 10 μM concentration. Antimycin-A (AA Blocks) stock solutions were maintained in DMSO at -20°C with a concentration of 5 mM. Rapamycin or Antimycin-A were added directly to cooled agar and plates were used immediately.

To assay growth differences on Rapamycin or Antimycin-A minimal media agar, fungal cells in the minimal media agar lawns were resuspended in PBS, washed, counted, and diluted to 1x10^6^ cells/mL. Cell suspensions were serially diluted 1:3 and spotted onto minimal media plates for 48 h at 30°C.

### Galleria mellonella infections

Final instar *G*. *mellonella* were obtained through Vanderhorst Wholesale, St. Marys, Ohio, USA. Healthy (motile, cream colored) larvae weighing approximately 175 to 225 mg were selected and left to acclimate overnight. *C. neoformans (*WT*, mfe2*Δ*, or pot1*Δ) cultures were grown in YPD overnight. Groups of 15 larvae were injected with 10 μl of PBS or 10 μl of washed *C. neoformans* cultures suspended in PBS at a density of 1x10^7^ cells/ml, which is an injected inoculum of 1x10^5^ cell/larva. *G*. *mellonella* larvae and pupae were kept at 30°C and monitored daily for survival for 10 days. Survival was assessed by movement upon stimulus with a pipette.

### Mouse model virulence studies

#### Murine Infection

Before the infection, *C. neoformans* yeast cells of WT (KN99α), *pot1*Δ on day 11, and *mfe2*Δ from liquid cultures were washed twice with PBS. A/J mice were put under anesthesia using 100 mg/kg of Ketamine (Covetrus, OH, US) and 10 mg/kg of Xylazine (Covetrus, OH, US), 100 µL of PBS containing 5×10^5^ cryptococcal cells were inoculated in the retro orbital vain. Survival curves were compared (n=7 mice/strain) post intravenous infection.

To compare colony forming units (CFU), n=3 mice/strain were euthanized on the same day for each strain (WT(KN99α) on day 5, *pot1*Δ on day 11, and *mfe2*Δ on day 19). The lungs and brain were aseptically removed, and transversal sections of the lung tissue were randomly collected for histological processing. The remaining tissue was weighed. The lungs were homogenized in 2 mL of PBS, and 100 μL were plated on glutamate-substituted minimal media agar (15.0 mM glucose, 10.0 mM MgSO_4_, 29.4 mM KH_2_PO_4_, 13.0 mM glutamate, 3.0 mM vitamin B_1_, pH 5.5 and 1% Pen/Strep). Plates were incubated at 30 °C (2 d for WT and pot1Δ and 4 d for mfe2Δ) and CFU were counted.

#### Histopathology

To evaluate the lung tissue after infection, lung transversal sections were fixed in 10% buffered formalin and embedded in paraffin using standard protocol. Histological slides were stained by the Johns Hopkins University Reference Histology Laboratory with hematoxylin and eosin (HE) for evaluation of the cellular structure of the organ, silver staining was performed to evaluate the presence of the fungal cell wall, and Masson staining to evaluate collagen deposition. Slides were scanned using the NanoZoomer S360MD Slide scanner (Hamamatsu Corporation - Bridgewater, NJ, US) by the he Johns Hopkins University Oncology Tissue and Imaging Service (OTIS) Core Laboratory (*Grant number, P30 CA006973)*.

#### Cytokine levels

For cytokine quantification, lung homogenates were aliquoted into microtubes with 500 μL of Roche cOmplete protein inhibitor (Roche - Indianapolis, IN, US). Cytokine analysis was performed by Enzyme-Linked Immunosorbent Assay (ELISA) using commercial kit (Thermo Fisher Scientific - Waltham, MA, US) for the following cytokines: IL1-β, IFN-γ, IL-4, IL-6 and IL-10, using the manufacturer’s instructions. IL-12p70 was measured using the BD OptEIA™ Mouse IL-12 p70 ELISA Set (Becton, Dickinson and Company Franklin Lakes, NJ, US), using the manufacturer’s instructions.

### TEM and peroxisomal measurements

Strains were grown for 2 h and 16 h at 30°C with a starting cell density of 5x10^6^ cells/mL in 5 mL media. To compare peroxisomal induction across medias, cell cultures were collected after 2 h of growth. The medias tested at 2 h were standard minimal media (glucose/glycine), acetate substituted minimal media (acetate/glycine), and glutamate substituted minimal media (glucose/glutamate). At 16 h, WT, *mfe2*Δ, and *pot1*Δ grown in minimal media (glucose/glycine) were fixed and stained to compare across time points. Peroxisomal measurements were performed blinded to the strain/media using the freehand measure tool on FIJI image processing software.

To confirm peroxisomal identification, 2 h minimal media samples were stained with resuspend dried rabbit anti-SKL serum in 100 microliters of distilled, deionized water. The rabbit anti-SKL was a kind gift from Dr. Richard Rachubinski. Staining was performed using Gary Eitzen’s immunoEM protocol. Use at 1:100 dilution for IF and at 1:1000 for western blot. (Data not shown here)

### qPCR

WT fungal cells were inoculated at a starting cell density of 5x10^6^ cells/mL in 5 mL media. Antimycin-A (AA Blocks) was added to cultures at 0 h at a concentration of 1 μM and 2.5 μM. Samples were spun down at 4°C at three time points: 0 h, 2 h, and 6 h post-inoculation and resuspended in Trizol at -80°C (ThermoFisher).

Primer sequences were designed using Primer Express. For all experiments, Sybr Green was used and transcripts were normalized to expression of actin. The following primer sequences were used: CNAG_07747 Pox1, acyl-CoA oxidase (F: GGGAGCCGCCATCAAAG, R: CCGATCGTTGAGGTATGCAA). CNAG_00490 Pot1 acetyl-CoA acyltransferase (F: GGCATCCGAGAAGGTGTGA, R: GCGAATGCGGGTTTGAGTT), CNAG_03019 Faa2, long-chain acyl-CoA synthetase (F: CCTGGTACTTGCGGTCGTTT R: TCGGGAACATCAACAAGTTTGA). CNAG_05721 MFE2 (F: TGAGATGGGAAAGATCGAAAGG R: CCGCAGAAGGAGTGAAAGTGT)

### mRNA-Seq comparisons

Raw fastq sequencing files from each publication were downloaded directly from its respective Gene Expression Omnibus database. Fastq files were aligned to the C. neoformans reference genome from EuPath FungiDB (H99 v63) using Salmon (1.10.2). The resulting transcript counts were analyzed via Bioconductor DESeq2 (1.48.1) in R (4.4.2). Significance was declared at a Benjamini-Hochberg adjusted p value of less than 0.05. Log2FC values from each individual dataset were then aligned according to gene to generate heatmap figures.

## References

1. Steenbergen, J.N., H.A. Shuman, and A. Casadevall, Cryptococcus neoformans Interactions with Amoebae Suggest an Explanation for Its Virulence and Intracellular Pathogenic Strategy in Macrophages. Proceedings of the National Academy of Sciences of the United States of America, 2001. 98(26): p. 15245–15250.

2. Abdul Rahman, N.S.N., N.W. Abdul Hamid, and K. Nadarajah, Effects of Abiotic Stress on Soil Microbiome. Int J Mol Sci, 2021. 22(16).

3. Serna-Espinosa, B.N., D. Guzmán-Sanabria, M. Forero-Castro, P. Escandón, and Z.A. Sánchez-Quitian, Environmental Status of Cryptococcus neoformans and Cryptococcus gattii in Colombia. J Fungi (Basel), 2021. 7(6).

4. Liu, T.B., D.S. Perlin, and C. Xue, Molecular mechanisms of cryptococcal meningitis. Virulence, 2012. 3(2): p. 173–81.

5. Hu, G., P.-Y. Cheng, A. Sham, J.R. Perfect, and J.W. Kronstad, Metabolic adaptation in Cryptococcus neoformans during early murine pulmonary infection. Molecular Microbiology, 2008. 69(6): p. 1456–1475.

6. Derengowski Lda, S., H.C. Paes, P. Albuquerque, A.H. Tavares, L. Fernandes, I. Silva-Pereira, and A. Casadevall, The transcriptional response of Cryptococcus neoformans to ingestion by Acanthamoeba castellanii and macrophages provides insights into the evolutionary adaptation to the mammalian host. Eukaryot Cell, 2013. 12(5): p. 761–74.

7. Fan, W., P.R. Kraus, M.J. Boily, and J. Heitman, Cryptococcus neoformans gene expression during murine macrophage infection. Eukaryot Cell, 2005. 4(8): p. 1420–33.

8. Steen, B.R., S. Zuyderduyn, D.L. Toffaletti, M. Marra, S.J. Jones, J.R. Perfect, and J. Kronstad, Cryptococcus neoformans gene expression during experimental cryptococcal meningitis. Eukaryot Cell, 2003. 2(6): p. 1336–49.

9. Rude, T.H., D.L. Toffaletti, G.M. Cox, and J.R. Perfect, Relationship of the glyoxylate pathway to the pathogenesis of Cryptococcus neoformans. Infect Immun, 2002. 70(10): p. 5684–94.

10. Idnurm, A., Y.-S. Bahn, K. Nielsen, X. Lin, J.A. Fraser, and J. Heitman, Deciphering the Model Pathogenic Fungus Cryptococcus Neoformans. Nature Reviews Microbiology, 2005. 3(10): p. 753–764.

11. Cramer Kari, L., D. Gerrald Quincy, B. Nichols Connie, S. Price Michael, and J.A. Alspaugh, Transcription Factor Nrg1 Mediates Capsule Formation, Stress Response, and Pathogenesis in Cryptococcus neoformans. Eukaryotic Cell, 2006. 5(7): p. 1147–1156.

12. Summers, D.K., D.S. Perry, B. Rao, and H.D. Madhani, Coordinate genomic association of transcription factors controlled by an imported quorum sensing peptide in Cryptococcus neoformans. PLOS Genetics, 2020. 16(9): p. e1008744.

13. Homer, C.M., D.K. Summers, A.I. Goranov, S.C. Clarke, D.L. Wiesner, J.K. Diedrich, J.J. Moresco, D. Toffaletti, R. Upadhya, I. Caradonna, S. Petnic, V. Pessino, C.A. Cuomo, J.K. Lodge, J. Perfect, J.R. Yates, 3rd, K. Nielsen, C.S. Craik, and H.D. Madhani, *Intracellular Action of a Secreted Peptide Required for Fungal Virulence*. Cell Host Microbe, 2016. 19(6): p. 849–64.

14. Tian, X., G.-J. He, P. Hu, L. Chen, C. Tao, Y.-L. Cui, L. Shen, W. Ke, H. Xu, Y. Zhao, Q. Xu, F. Bai, B. Wu, E. Yang, X. Lin, and L. Wang, Cryptococcus neoformans sexual reproduction is controlled by a quorum sensing peptide. Nature Microbiology, 2018. 3(6): p. 698–707.

15. Chang, A.L. and T.L. Doering, Maintenance of Mitochondrial Morphology in Cryptococcus neoformans Is Critical for Stress Resistance and Virulence. mBio, 2018. 9(6): p. 10.1128/mbio.01375-18.

16. Caza, M., G. Hu, M. Price, J.R. Perfect, and J.W. Kronstad, The Zinc Finger Protein Mig1 Regulates Mitochondrial Function and Azole Drug Susceptibility in the Pathogenic Fungus Cryptococcus neoformans. mSphere, 2016. 1(1).

17. Caza, M., G. Hu, E.D. Nielson, M. Cho, W.H. Jung, and J.W. Kronstad, The Sec1/Munc18 (SM) protein Vps45 is involved in iron uptake, mitochondrial function and virulence in the pathogenic fungus Cryptococcus neoformans. PLoS Pathog, 2018. 14(8): p. e1007220.

18. Xu, S.C., M.D. He, Y.H. Lu, L. Li, M. Zhong, Y.W. Zhang, Y. Wang, Z.P. Yu, and Z. Zhou, Nickel exposure induces oxidative damage to mitochondrial DNA in Neuro2a cells: the neuroprotective roles of melatonin. J Pineal Res, 2011. 51(4): p. 426–33.

19. Kretschmer, M., J. Wang, and J.W. Kronstad, *Peroxisomal and mitochondrial* β*-oxidation pathways influence the virulence of the pathogenic fungus Cryptococcus neoformans*. Eukaryot Cell, 2012. 11(8): p. 1042–54.

20. Vasconcelles, M.J., Y. Jiang, K. McDaid, L. Gilooly, S. Wretzel, D.L. Porter, C.E. Martin, and M.A. Goldberg, Identification and characterization of a low oxygen response element involved in the hypoxic induction of a family of Saccharomyces cerevisiae genes. Implications for the conservation of oxygen sensing in eukaryotes. J Biol Chem, 2001. 276(17): p. 14374–84.

21. Kwast, K.E., P.V. Burke, B.T. Staahl, and R.O. Poyton, Oxygen sensing in yeast: evidence for the involvement of the respiratory chain in regulating the transcription of a subset of hypoxic genes. Proc Natl Acad Sci U S A, 1999. 96(10): p. 5446–51.

22. Gleason, J.E., D.J. Corrigan, J.E. Cox, A.R. Reddi, L.A. McGinnis, and V.C. Culotta, Analysis of Hypoxia and Hypoxia-Like States through Metabolite Profiling. PLOS ONE, 2011. 6(9): p. e24741.

23. Lee, H., C.M. Bien, A.L. Hughes, P.J. Espenshade, K.J. Kwon Chung, and Y.C. Chang, Cobalt chloride, a hypoxia mimicking agent, targets sterol synthesis in the pathogenic fungus Cryptococcus neoformans. Molecular microbiology, 2007. 65(4): p. 1018–1033.

24. Hommel, B., A. Sturny-Leclère, S. Volant, N. Veluppillai, M. Duchateau, C.-H. Yu, V. Hourdel, H. Varet, M. Matondo, J.R. Perfect, A. Casadevall, F. Dromer, and A. Alanio, Cryptococcus neoformans resists to drastic conditions by switching to viable but non-culturable cell phenotype. PLOS Pathogens, 2019. 15(7): p. e1007945.

25. Shani, N., P.A. Watkins, and D. Valle, PXA1, a possible Saccharomyces cerevisiae ortholog of the human adrenoleukodystrophy gene. Proc Natl Acad Sci U S A, 1995. 92(13): p. 6012–6.

26. Hettema, E.H., C.W. van Roermund, B. Distel, M. van den Berg, C. Vilela, C. Rodrigues-Pousada, R.J. Wanders, and H.F. Tabak, The ABC transporter proteins Pat1 and Pat2 are required for import of long-chain fatty acids into peroxisomes of Saccharomyces cerevisiae. Embo j, 1996. 15(15): p. 3813–22.

27. van Roermund, C.W., Y. Elgersma, N. Singh, R.J. Wanders, and H.F. Tabak, The membrane of peroxisomes in Saccharomyces cerevisiae is impermeable to NAD(H) and acetyl-CoA under in vivo conditions. Embo j, 1995. 14(14): p. 3480–6.

28. Knoll, L.J., D.R. Johnson, and J.I. Gordon, Biochemical studies of three Saccharomyces cerevisiae acyl-CoA synthetases, Faa1p, Faa2p, and Faa3p. J Biol Chem, 1994. 269(23): p. 16348-56.

29. Wang, T., Y. Luo, and G.M. Small, The POX1 gene encoding peroxisomal acyl-CoA oxidase in Saccharomyces cerevisiae is under the control of multiple regulatory elements. Journal of Biological Chemistry, 1994. 269(39): p. 24480–24485.

30. Gabriel, F., I. Accoceberry, J.J. Bessoule, B. Salin, M. Lucas-Guérin, S. Manon, K. Dementhon, and T. Noël, A Fox2-dependent fatty acid ß-oxidation pathway coexists both in peroxisomes and mitochondria of the ascomycete yeast Candida lusitaniae. PLoS One, 2014. 9(12): p. e114531.

31. Igual, J.C., C. González-Bosch, L. Franco, and J.E. Pérez-Ortín, The POT1 gene for yeast peroxisomal thiolase is subject to three different mechanisms of regulation. Mol Microbiol, 1992. 6(14): p. 1867–75.

32. Cheong, J.W. and J. McCormack, Fluconazole resistance in cryptococcal disease: emerging or intrinsic? Med Mycol, 2013. 51(3): p. 261–9.

33. Mpoza, E., J. Rhein, and M. Abassi, Emerging fluconazole resistance: Implications for the management of cryptococcal meningitis. Med Mycol Case Rep, 2018. 19: p. 30–32.

34. Chelstowska, A., Y. Jia, B. Rothermel, and R.A. Butow, Retrograde regulation: a novel path of communication between mitochondria, the nucleus, and peroxisomes in yeast. Canadian journal of botany, 1995. 73(S1): p. 205–207.

35. Liu, Z. and R.A. Butow, Mitochondrial Retrograde Signaling. Annual Review of Genetics, 2006. 40(Volume 40, 2006): p. 159–185.

36. Jung, K.-W., D.-H. Yang, S. Maeng, K.-T. Lee, Y.-S. So, J. Hong, J. Choi, H.-J. Byun, H. Kim, S. Bang, M.-H. Song, J.-W. Lee, M.S. Kim, S.-Y. Kim, J.-H. Ji, G. Park, H. Kwon, S. Cha, G.L. Meyers, L.L. Wang, J. Jang, G. Janbon, G. Adedoyin, T. Kim, A.K. Averette, J. Heitman, E. Cheong, Y.-H. Lee, Y.-W. Lee, and Y.-S. Bahn, Systematic functional profiling of transcription factor networks in Cryptococcus neoformans. Nature Communications, 2015. 6(1): p. 6757.

37. Liu, O.W., C.D. Chun, E.D. Chow, C. Chen, H.D. Madhani, and S.M. Noble, Systematic Genetic Analysis of Virulence in the Human Fungal Pathogen Cryptococcus neoformans. Cell, 2008. 135(1): p. 174–188.

38. Yu, C.H., Y. Chen, C.A. Desjardins, J.L. Tenor, D.L. Toffaletti, C. Giamberardino, A. Litvintseva, J.R. Perfect, and C.A. Cuomo, Landscape of gene expression variation of natural isolates of Cryptococcus neoformans in response to biologically relevant stresses. Microb Genom, 2020. 6(1).

39. Tate, J.J. and T.G. Cooper, Tor1/2 Regulation of Retrograde Gene Expression in Saccharomyces cerevisiae Derives Indirectly as a Consequence of Alterations in Ammonia Metabolism*. Journal of Biological Chemistry, 2003. 278(38): p. 36924–36933.

40. Stolp, Z.D., M. Kulkarni, Y. Liu, C. Zhu, A. Jalisi, S. Lin, A. Casadevall, K.W. Cunningham, F.J. Pineda, X. Teng, and J.M. Hardwick, Yeast cell death pathwayrequiring AP-3 vesicle trafficking leads to vacuole/lysosome membrane permeabilization. Cell Reports, 2022. 39(2): p. 110647.

41. Liao, X. and R.A. Butow, RTG1 and RTG2: Two yeast genes required for a novel path of communication from mitochondria to the nucleus. Cell, 1993. 72(1): p. 61–71.

42. Liew, K.L., J.M. Jee, I. Yap, and P.V.C. Yong, In Vitro Analysis of Metabolites Secreted during Infection of Lung Epithelial Cells by Cryptococcus neoformans. PLOS ONE, 2016. 11(4): p. e0153356.

43. Baker, R.P. and A. Casadevall, Reciprocal modulation of ammonia and melanin production has implications for cryptococcal virulence. Nature Communications, 2023. 14(1): p. 849.

44. Fu, M.S., C. Coelho, C.M. De Leon-Rodriguez, D.C.P. Rossi, E. Camacho, E.H. Jung, M. Kulkarni, and A. Casadevall, Cryptococcus neoformans urease affects the outcome of intracellular pathogenesis by modulating phagolysosomal pH. PLOS Pathogens, 2018. 14(6): p. e1007144.

45. Cox, G.M., J. Mukherjee, G.T. Cole, A. Casadevall, and J.R. Perfect, Urease as a virulence factor in experimental cryptococcosis. Infect Immun, 2000. 68(2): p. 443–8.

46. Croft, B., G.R. Wentworth, R.V. Martin, W.R. Leaitch, J.G. Murphy, B.N. Murphy, J.K. Kodros, J.P.D. Abbatt, and J.R. Pierce, Contribution of Arctic seabird-colony ammonia to atmospheric particles and cloud-albedo radiative effect. Nature Communications, 2016. 7(1): p. 13444.

47. Komeili, A., K.P. Wedaman, E.K. O’Shea, and T. Powers, Mechanism of metabolic control. Target of rapamycin signaling links nitrogen quality to the activity of the Rtg1 and Rtg3 transcription factors. J Cell Biol, 2000. 151(4): p. 863–78.

48. Dilova, I., C.-Y. Chen, and T. Powers, Mks1 in Concert with TOR Signaling Negatively Regulates RTG Target Gene Expression in S. cerevisiae. Current Biology, 2002. 12(5): p. 389–395.

49. Albuquerque, P., A.M. Nicola, E. Nieves, H.C. Paes, P.R. Williamson, I. Silva-Pereira, and A. Casadevall, *Quorum Sensing-Mediated,* Cell Density-Dependent Regulation of Growth and Virulence in Cryptococcus neoformans. mBio, 2014. 5(1): p. 10.1128/mbio.00986-13.

50. Hommel, B., L. Mukaremera, R.J.B. Cordero, C. Coelho, C.A. Desjardins, A. Sturny-Leclère, G. Janbon, J.R. Perfect, J.A. Fraser, A. Casadevall, C.A. Cuomo, F. Dromer, K. Nielsen, and A. Alanio, Titan cells formation in Cryptococcus neoformans is finely tuned by environmental conditions and modulated by positive and negative genetic regulators. PLOS Pathogens, 2018. 14(5): p. e1006982.

51. Smith, J.J., T.W. Brown, G.A. Eitzen, and R.A. Rachubinski, Regulation of peroxisome size and number by fatty acid beta -oxidation in the yeast yarrowia lipolytica. J Biol Chem, 2000. 275(26): p. 20168–78.

52. Piekarska, K., G. Hardy, E. Mol, J. van den Burg, K. Strijbis, C. van Roermund, M. van den Berg, and B. Distel, *The activity of the glyoxylate cycle in peroxisomes of Candida albicans depends on a functional* β*-oxidation pathway: evidence for reduced metabolite transport across the peroxisomal membrane*. Microbiology, 2008. 154(10): p. 3061–3072.

53. Ingavale, S.S., Y.C. Chang, H. Lee, C.M. McClelland, M.L. Leong, and K.J. Kwon-Chung, Importance of mitochondria in survival of Cryptococcus neoformans under low oxygen conditions and tolerance to cobalt chloride. PLoS Pathog, 2008. 4(9): p. e1000155.

54. Lindegren, C.C., S. Nagai, and H. Nagai, Induction of Respiratory Deficiency in Yeast by Manganese, Copper, Cobalt and Nickel. Nature, 1958. 182(4633): p. 446–448.

55. Parikh, V.S., M.M. Morgan, R. Scott, L.S. Clements, and R.A. Butow, The mitochondrial genotype can influence nuclear gene expression in yeast. Science, 1987. 235(4788): p. 576–80.

56. Crespo, J.L., T. Powers, B. Fowler, and M.N. Hall, The TOR-controlled transcription activators GLN3, RTG1, and RTG3 are regulated in response to intracellular levels of glutamine. Proceedings of the National Academy of Sciences, 2002. 99(10): p. 6784–6789.

57. Bui, T.H.D. and K. Labedzka-Dmoch, RetroGREAT signaling: The lessons we learn from yeast. IUBMB Life, 2024. 76(1): p. 26–37.

58. Chelstowska, A. and R.A. Butow, RTG Genes in Yeast That Function in Communication between Mitochondria and the Nucleus Are Also Required for Expression of Genes Encoding Peroxisomal Proteins *. Journal of Biological Chemistry, 1995. 270(30): p. 18141–18146.

59. Jazwinski, S.M. and A. Kriete, The yeast retrograde response as a model of intracellular signaling of mitochondrial dysfunction. Front Physiol, 2012. 3: p. 139.

60. Moreno-Velásquez, S.D., S.H. Tint, V. Del Olmo Toledo, S. Torsin, S. De, and J.C. Pérez, The Regulatory Proteins Rtg1/3 Govern Sphingolipid Homeostasis in the Human-Associated Yeast Candida albicans. Cell Rep, 2020. 30(3): p. 620–629.e6.

61. Liu, Z., P. Basso, S. Hossain, S.D. Liston, N. Robbins, L. Whitesell, S.M. Noble, and L.E. Cowen, Multifactor transcriptional control of alternative oxidase induction integrates diverse environmental inputs to enable fungal virulence. Nature Communications, 2023. 14(1): p. 4528.

62. Lazazzera, B.A., Quorum sensing and starvation: signals for entry into stationary phase. Curr Opin Microbiol, 2000. 3(2): p. 177–82.

63. Lee, H., Y.C. Chang, G. Nardone, and K.J. Kwon-Chung, TUP1 disruption in Cryptococcus neoformans uncovers a peptide-mediated density-dependent growth phenomenon that mimics quorum sensing. Molecular Microbiology, 2007. 64(3): p. 591–601.

64. Hashim, Z., Y. Mukai, T. Bamba, and E. Fukusaki, Metabolic profiling of retrograde pathway transcription factors rtg1 and rtg3 knockout yeast. Metabolites, 2014. 4(3): p. 580–98.

65. Janssen, J.J.E., S. Grefte, J. Keijer, and V.C.J. de Boer, Mito-Nuclear Communication by Mitochondrial Metabolites and Its Regulation by B-Vitamins. Front Physiol, 2019. 10: p. 78.

66. Leonardi, R., Y.M. Zhang, C.O. Rock, and S. Jackowski, Coenzyme A: back in action. Prog Lipid Res, 2005. 44(2-3): p. 125–53.

67. Silva Vanessa, K.A., S. Bhattacharya, K. Oliveira Natalia, G. Savitt Anne, D. Zamith-Miranda, D. Nosanchuk Joshua, and C. Fries Bettina, Replicative Aging Remodels the Cell Wall and Is Associated with Increased Intracellular Trafficking in Human Pathogenic Yeasts. mBio, 2022. 13(1): p. e00190–22.

68. Homann, O.R., J. Dea, S.M. Noble, and A.D. Johnson, A phenotypic profile of the Candida albicans regulatory network. PLoS genetics, 2009. 5(12): p. e1000783.

69. Ramírez, M.A. and M.C. Lorenz, Mutations in alternative carbon utilization pathways in Candida albicans attenuate virulence and confer pleiotropic phenotypes. Eukaryot Cell, 2007. 6(2): p. 280–90.

70. Otzen, C., B. Bardl, I.D. Jacobsen, M. Nett, and M. Brock, *Candida albicans Utilizes a Modified* β*-Oxidation Pathway for the Degradation of Toxic Propionyl-CoA\**. Journal of Biological Chemistry, 2014. 289(12): p. 8151–8169.

71. Loftus, B.J., E. Fung, P. Roncaglia, D. Rowley, P. Amedeo, D. Bruno, J. Vamathevan, M. Miranda, I.J. Anderson, J.A. Fraser, J.E. Allen, I.E. Bosdet, M.R. Brent, R. Chiu, T.L. Doering, M.J. Donlin, C.A. D’Souza, D.S. Fox, V. Grinberg, J. Fu, M. Fukushima, B.J. Haas, J.C. Huang, G. Janbon, S.J. Jones, H.L. Koo, M.I. Krzywinski, J.K. Kwon-Chung, K.B. Lengeler, R. Maiti, M.A. Marra, R.E. Marra, C.A. Mathewson, T.G. Mitchell, M. Pertea, F.R. Riggs, S.L. Salzberg, J.E. Schein, A. Shvartsbeyn, H. Shin, M. Shumway, C.A. Specht, B.B. Suh, A. Tenney, T.R. Utterback, B.L. Wickes, J.R. Wortman, N.H. Wye, J.W. Kronstad, J.K. Lodge, J. Heitman, R.W. Davis, C.M. Fraser, and R.W. Hyman, The genome of the basidiomycetous yeast and human pathogen Cryptococcus neoformans. Science, 2005. 307(5713): p. 1321–4.

72. Stolp, Z.D., M. Kulkarni, Y. Liu, C. Zhu, A. Jalisi, S. Lin, A. Casadevall, K.W. Cunningham, F.J. Pineda, X. Teng, and J.M. Hardwick, Yeast cell death pathway requiring AP-3 vesicle trafficking leads to vacuole/lysosome membrane permeabilization. Cell Rep, 2022. 39(2): p. 110647.

