## Supplementary material for "Peroxisomes regulate virulence and cell density sensing in *Cryptococcus neoformans*": Supp Fig.

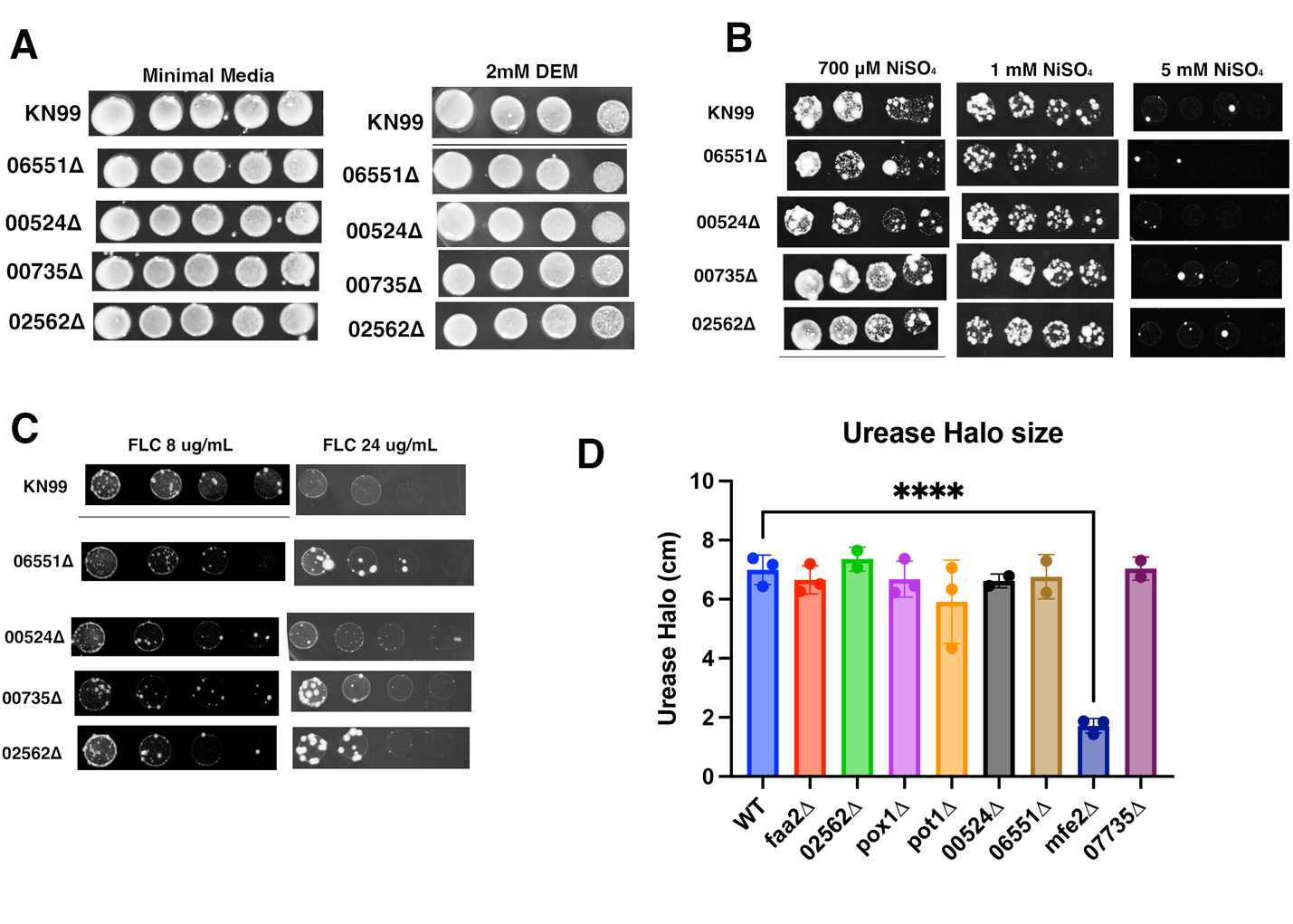

**Supplemental Figure 1: Screening of additional candidate genes upregulated during nickel exposure.**

**A)** Exposure to oxidative stress inducer DEM.

**B)** Exposure to nickel sulfate.

**C)** Exposure to fluconazole.

**D)** Urease halo size.

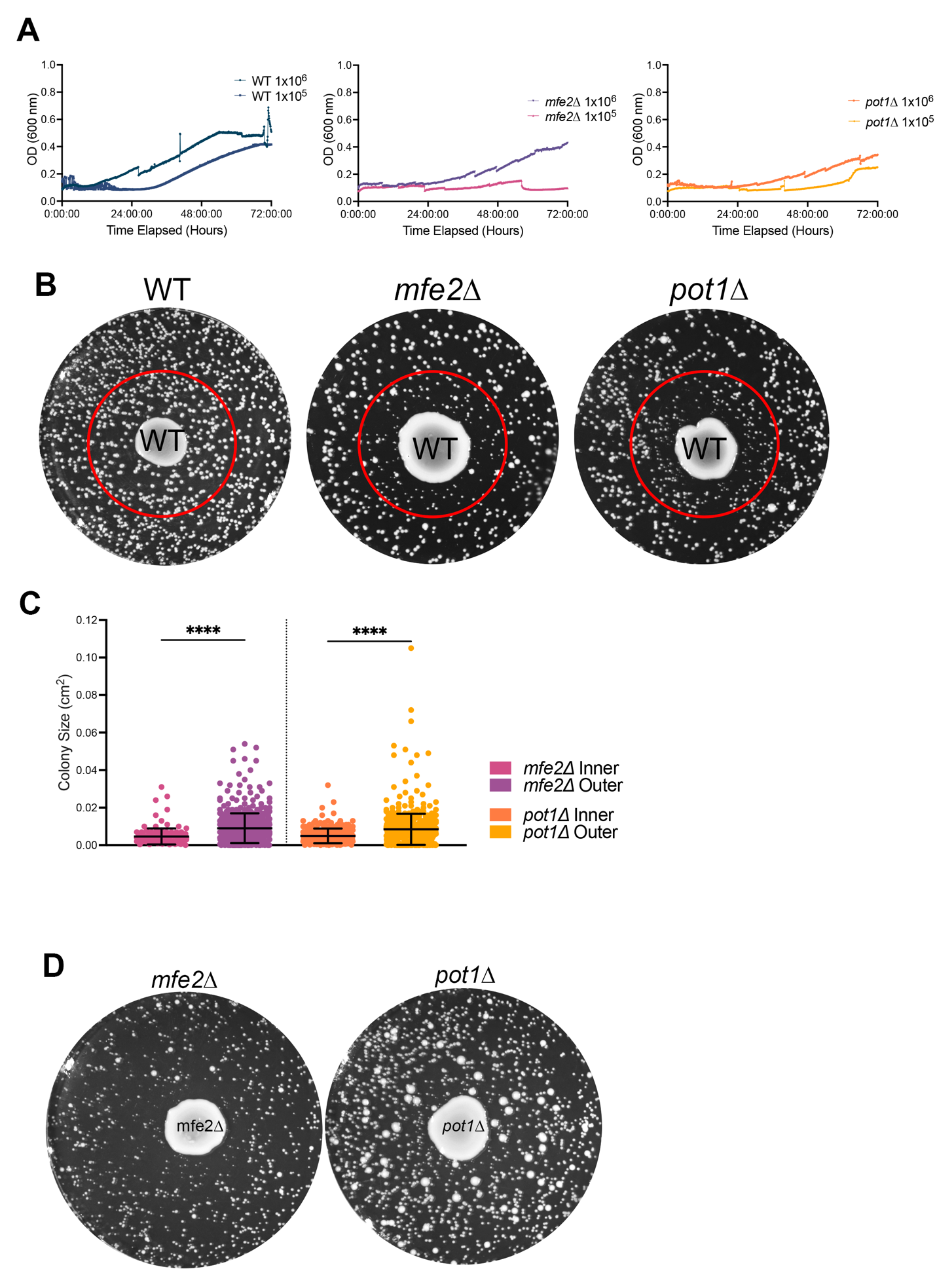

**Supplemental Figure 2: Intermediate growth phenotypes and zone of inhibition.**

**A)** Intermediate growth inhibition phenotypes occur at the cell densities initially used to test virulence factors. Growth curves (O.D. 600 nm) comparing cell densities 1x10^6^ cells/mL and 1x10^5^ cells/mL in minimal media over 72 h.

**B)** Zone of mutant colony growth inhibition (red circle) surrounding the 100μL WT rescue colony.

**C)** Comparison of colony size in the zone of inhibition (red circle) to the peripheral colonies. (Kruskall-Wallis with Dunn’s multiple comparisons test. ****, p<0.001)

**D)** The zone of growth inhibition also occurs when the 100 μL rescue colony is the mutant strain instead of WT.

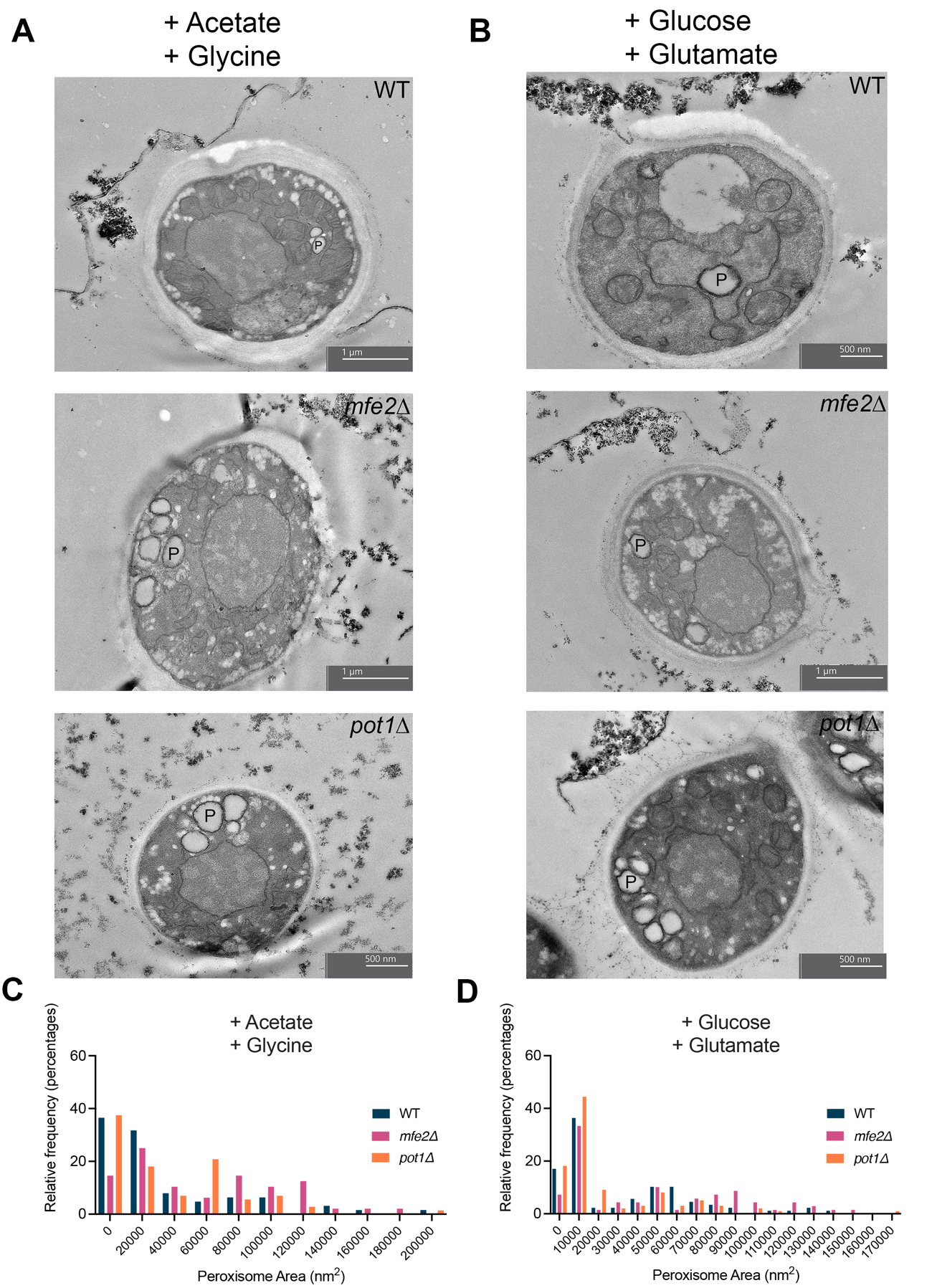

**Supplemental Figure 3: Initial peroxisome induction occurs regardless of media.**

**A)** Representative TEM images of WT, *mfe2∆*, and *pot1∆* after 2 h of growth in Acetate substituted minimal media. P indicates peroxisome.

**B)** Representative TEM images of WT, *mfe2∆*, and *pot1∆* after 2 h of growth in Glutamate substituted minimal media. P indicates peroxisome.

**C)** Peroxisome area size distribution after 2 h of growth in Acetate substituted minimal media.

**D)** Peroxisome area size distribution after 2 h of growth in Glutamate substituted minimal media.

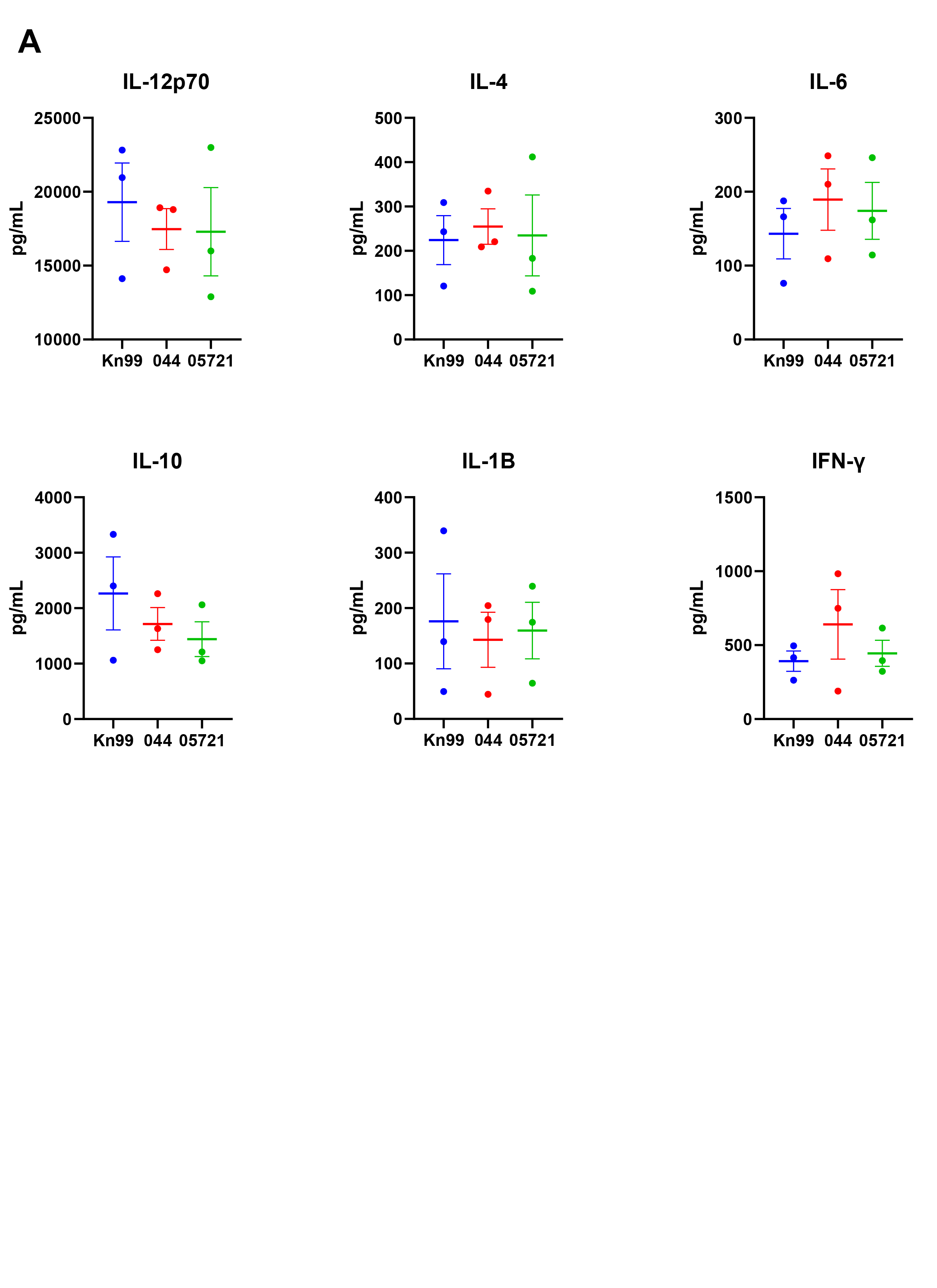

**Supplemental Figure 4: Lung cytokine levels measured by ELISA.**

**A)** Cytokine profiles of WT (blue), *pot1∆* (red), and *mfe2∆* (green).
